# The impact of incomplete taxon sampling on inference of gene flow by Bayesian and summary methods using genomic sequence data

**DOI:** 10.1101/2025.11.05.686701

**Authors:** Sirui Cheng, Thomas Flouri, Tianqi Zhu, Ziheng Yang

**Affiliations:** Department of Genetics, Evolution, and Environment, University College London, Gower Street, London WC1E 6BT, UK; State Key Laboratory of Mathematical Science, Academy of Mathematics and Systems Science, Chinese Academy of Sciences, Beijing 100190, China; University of Chinese Academy of Sciences, Beijing 100049, China

**Keywords:** Bayesian test, BPP, ghost introgression, introgression, migration, multispecies coalescent, Savage-Dickey, taxon sampling

## Abstract

Interspecific gene flow is commonly inferred using genomic data under the multispecies coalescent (MSC) model. The rate of gene flow is measured by the expected proportion of immigrants in the recipient population at the time of hybridization/introgression. Incomplete taxon sampling can impact inference of gene flow in multiple ways. First unsampled ghost lineages that are sources of introgression may mislead inference of gene flow in analysis of genomic data from modern species. Second incomplete taxon sampling causes merges of branches on the species phylogeny, which represent populations of different sizes, and complicates the definition and estimation of the introgression probability. We use mathematical analysis and computer simulation to examine the impact of incomplete taxon sampling on inference of gene flow and estimation of its rate using genomic data. We introduce a Bayesian testing approach to select models of gene flow for a species triplet (such as inflow, outflow, and ghost introgression), using the Savage-Dickey density ratio to calculate Bayes factors. We show that the approach has excellent power and specificity. We find that genomic data allow reliable estimation of the proportion of immigrants (rather than the number of immigrants), even when the assumed demographic model is incorrect due to incomplete taxon sampling. When population size differs among species, assuming the same size may lead to seriously biased estimates of the rate of gene flow. The ***f*** -branch approach is found to be effective in reducing the number of significant gene-flow events from triplet analyses, providing useful hypotheses for rigorous testing, but often to produce underestimates of the rate of gene flow. Our results highlight the need for improving summary methods to accommodate different population sizes and to infer gene flow between sister lineages.

## Introduction

Genomic sequences sampled from extant species provide a rich source of information concerning historical gene flow between species. Indeed gene flow has been detected using genomic data from a variety of species including animals as well as plants (Harrison and Larson, 2014; Edelman and Mallet, 2021). Gene flow has most often been inferred using simple summary methods that operate on species triplets (or quartets if an outgroup is included). They make use of either genomewide site-pattern counts, as in the case of the *D*-statistic or ABBA-BABA test (Green *et al*., 2010) and HyDe (Blischak *et al*., 2018), or reconstructed gene tree topologies, as in SNaQ (Solis-Lemus and Ane, 2016) and PhyloNet/MPL (Yu and Nakhleh, 2015). There is an issue of incomplete taxon sampling as the analysis uses only three species/populations, with one sequence sampled per species, even if genomic data are available from many species and multiple samples per species. Another scenario arises when there exist ghost lineages, i.e., extinct or unsampled lineages that contributed genetic materials to modern sampled species or their ancestors. In this study we use the term incomplete taxon sampling to refer to both scenarios.

Incomplete taxon sampling may influence our inference of interspecific gene flow in multiple ways. First, the presence of ghost lineages may mislead our inference of gene flow in analysis of sampled species (Beerli, 2004; Ottenburghs, 2020). Consider the phylogeny for six species with gene flow of Figure 1**a**. In analysis of the triplet *ACD*, species *E* and *F* are ghost lineages (which contribute genes to the sampled species *D*). Through simulations Tricou *et al*. (2022) found that introgression from an outgroup ghost lineage can cause the *D*-statistic to mis-identify the donor and the recipient populations involved in introgression. Huang *et al*. (2022, Figs. 6, 8) found that the absence of an intermediate ghost species does not have a great impact on BPP inference of gene flow, which detects ‘indirect’ gene flow via unsampled intermediate species as well as direct gene flow. Recently Pang and Zhang (2024) made the disturbing finding that commonly used triplet methods such as the *D*-statistic or ABBA-BABA test (Green *et al*., 2010), HyDe(Blischak *et al*., 2018), SNaQ (Solis-Lemus and Ane, 2016), and Phylonet/MPL (Yu and Nakhleh, 2015) cannot distinguish among different scenarios of introgression on a triplet tree, such as introgression from an outgroup ghost species, and inflow or outflow between non-sister ingroup species (Fig. 2**a**–**c**).

**Fig. 1.**
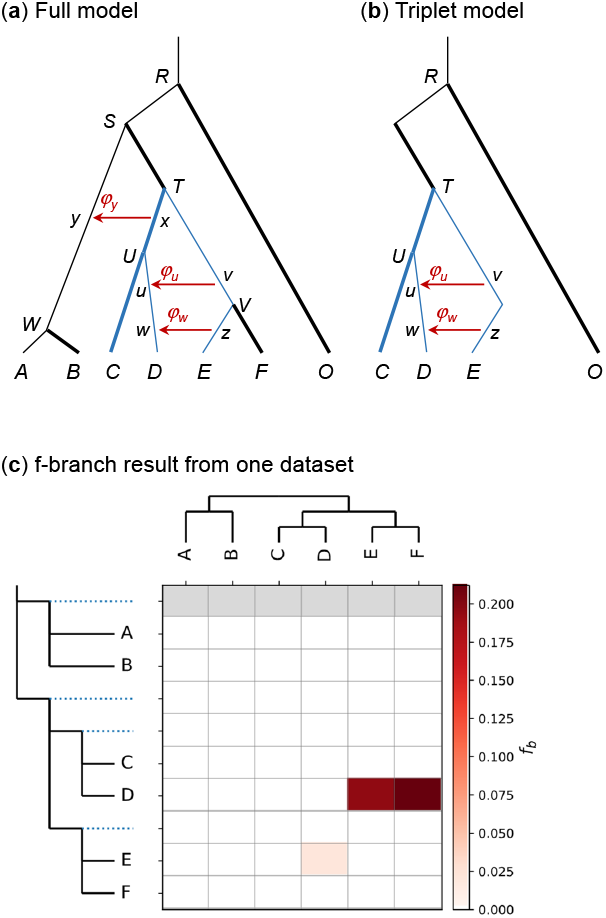
(**a**) Phylogeny for six species (*A*–*F*) plus an outgroup (*O*) with three introgression events used to simulate data. Population size was assumed to be either equal for all branches (*θ*_0_ = *θ*_1_ = 0.001) or different for thin and thick branches (*θ*_0_ = 0.001 and *θ*_1_ = 0.01, respectively). (**b**) Triplet species tree for the *CDE* triplet (with outgroup *O*) after other species in the tree of (**a**) are pruned off. Branches *TV* and *V E* in the original tree are merged into one branch *T E* in the triplet tree. (**c**) Heatmap of introgression probabilities produced in the *f* -branch analysis of a replicate dataset simulated under the model of **a**, with *L* = 1000 loci (with *S* = 4 sequences per locus and *n* = 500 sites per sequence). Three introgression events were inferred: *f*_*D*_ (*F*) = 0.212, *f*_*D*_ (*E*) = 0.194, and *f*_*E*_ (*D*) = 0.0194.

**Fig. 2.**
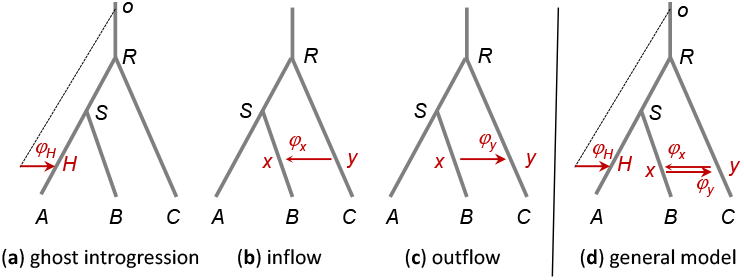
Three MSC-I models used to simulate data: (**a**) ghost introgression, (**b**) inflow from *C* → *B* (or *y* → *x*), and (**c**) outflow from *B* → *C* (or *x* → *y*). The data are then analyzed using bpp under the full model (**d**) to select the models of panels **a**–**c**. The data are also analyzed using summary methods HyDe and SNaQ, by including one (for HyDMe) or two (for SNaQ) distant outgroup species.

Second, multiple branches on the phylogeny, which correspond to species with different population sizes, may be merged into one branch in the triplet tree and assigned one population size parameter. The rate of gene flow is typically measured by the introgression probability, *φ*_*xy*_ — also denoted *γ* in HyDe (Blischak *et al*., 2018), SNaQ (Solis-Lemus and Ane, 2016) and PhyloNet/MPL (Yu and Nakhleh, 2015), and *f* in ADMIXTOOLS (Patterson *et al*., 2012) — and defined as the probability of immigrants in the recipient population *y* from the donor population *x* at the time of hybridization/introgression. The apparent changes in the size of the recipient population due to exclusion of species on the phylogeny may thus affect our definition and estimation of the rate of gene flow. For example, is it the proportion or the number of immigrants that affect the genomic data and can thus be reliably estimated using genomic data? Because the introgression probability *φ*_*xy*_ is also the probability that a sequence in the recipient population *y* is traced to the donor population *x* when one traces the genealogical history of the sample of sequences backwards in time, and because sequence data reflect information in the gene genealogies, one may expect inference using sequence data to be dominated by the expected proportion of immigrants (*φ*_*xy*_), rather than the expected number of immigrants (i.e., *N*_*y*_*φ*_*xy*_). This expectation is yet to be confirmed.

Third, branches in the phylogeny that are merged in the triplet tree may be involved in introgression in different directions at different time points, and those events are all merged into one introgression event with a single introgression probability, the expected value of which is poorly understood. For example, when we analyze the *CDE* triplet (Fig. 1**b**), the two introgression events, *v* → *u* and *z* → *w*, are merged into one event (from *E* to *D*). Do we expect the estimated introgression probability *φ*_*E*→*D*_ to be *φ*_*vu*_ + *φ*_*zw*_? What if the introgressions are in opposite directions (*u* → *v* and *z* → *w*, say)?

Fourth, one gene flow event involving ancestral branches on the phylogeny may appear in many triplets, and it may be challenging to combine many triplet estimates into one introgression probability for the shared gene-flow event. For example, the *x* → *y* introgression on the species tree of Figure 1**a** may show up in eight triplets, 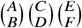, generated by selecting one of *A* and *B*, one of *C* and *D*, and one of *E* and *F*, and the eight estimates may differ from the true rate, affected by different population sizes on the triplet trees and by random sampling errors.

Indeed the *D*-statistic has been observed to detect gene flow in overwhelmingly many triplets in analyses of empirical data (e.g., Malinsky *et al*., 2018). The challenge of interpreting such triplet results motivated the development of the *f* -branch approach, which assembles results from the triplet analyses (Malinsky *et al*., 2018, 2021). The method assumes a binary species tree and attempts to move gene-flow events onto ancestral branches which may explain the detection of gene flow in many triplets. Ideally one would like each triplet analysis to reflect the true model, and the assembled results from all triplets to recover the true model for the full phylogeny. Currently it is unknown to what extent this ideal is fulfilled as we lack an understanding of the behavior of triplet methods when the three species are a subset of many species (as in the case of the *CDE* triplet of Fig. 1**b** when the true model is the one for six species of Fig. 1**a**). The inability of triplet methods to distinguish among competing models of gene flow as demonstrated by Pang and Zhang (2024) raises concerns about the *f* -branch approach, which relies on those methods.

Fifth, incomplete taxon sampling or taxon exclusion in triplet methods may seriously reduce the information content in the data and even cause unidentifiability issues. A gene flow event between non-sister lineages on the full phylogeny may become an event between sister lineages on the triplet tree and thus unidentifiable by summary methods. For example, the *v* → *u* and *z* → *w* gene flow in Fig. 1**a** is between nonsister species but becomes between sisters in the *ADE* triplet. Similarly use of a single sequence per species may create unidentifiability problems (Degnan, 2018; Jiao *et al*., 2021; Yang and Flouri, 2022). For example, the direction of gene flow between sister lineages is unidentifiable when only one sequence is sampled from each species, but is identifiable when multiple samples per species are used (Yang and Flouri, 2022, Fig. 10). Even when the model is identifiable, including multiple samples per species (in particular, from the recipient population) may boost the information concerning gene flow significantly.

In this study we consider both the presence of ghost lineages and the exclusion of lineages due to the restrictions of the triplet methods as instances of the general problem of incomplete taxon sampling, and study its impacts on inference of gene flow. Besides summary methods, we use the Bayesian method as implemented in the program BPP (Flouri *et al*., 2018, 2020, 2023). This includes efficient implementations of two classes of models of gene flow in the multispecies coalescent (MSC) framework: the MSC-introgression (MSC-I) model which assumes discrete gene flow with hybridization/introgression at a time point in the past (Wen and Nakhleh, 2018; Zhang *et al*., 2018; Flouri *et al*., 2020) and the MSC-migration (MSC-M) model which assumes continuous gene flow over an extended time period (Nielsen and Wakeley, 2001; Hey *et al*., 2018; Flouri *et al*., 2023).

Our paper is structured as follows. First we consider inference of ghost introgression, and describe a Bayesian testing approach to comparing models of gene flow in the case of species triplet, including ghost introgression, inflow, and outflow (Fig. 2**a**-**c**). This is inspired by Pang and Zhang (2024, Fig. 3), who demonstrated that commonly used summary triplet methods are unable to identify those models (see also Tricou *et al*. 2022). We use simulation to assess the performance of the Bayesian approach, in comparison with summary triplet methods such as *D* (Green *et al*., 2010), HyDe (Blischak *et al*., 2018), SNaQ (Solis-Lemus and Ane, 2016) and PhyloNet/MPL (Yu and Nakhleh, 2015). Second, we assess the impacts of the changing population size on the species tree due to incomplete taxon sampling on inference of gene flow. We study the estimation of the population size parameter *θ* for one population when *θ* actually changes over time. We conduct an asymptotic analysis considering the estimate using only two sequences per locus when the number of loci approaches infinity, and use simulation to study the case of multiple sequences per locus. We then study BPP estimation of the introgression probability between two sister species, and the performance of both BPP and summary methods in triplet data. We simulate data on a model species tree for six species to compare BPP and triplet methods for estimating the rate of gene flow and evaluate the *f* branch approach for summarizing triplet results. Overall our analyses suggest that assuming the same population size for all species when population size differs on the species tree may lead to serious biases in the estimated rate of gene flow, but the Bayesian method may provide reliable rate estimates even if taxon sampling is incomplete.

**Fig. 3.**
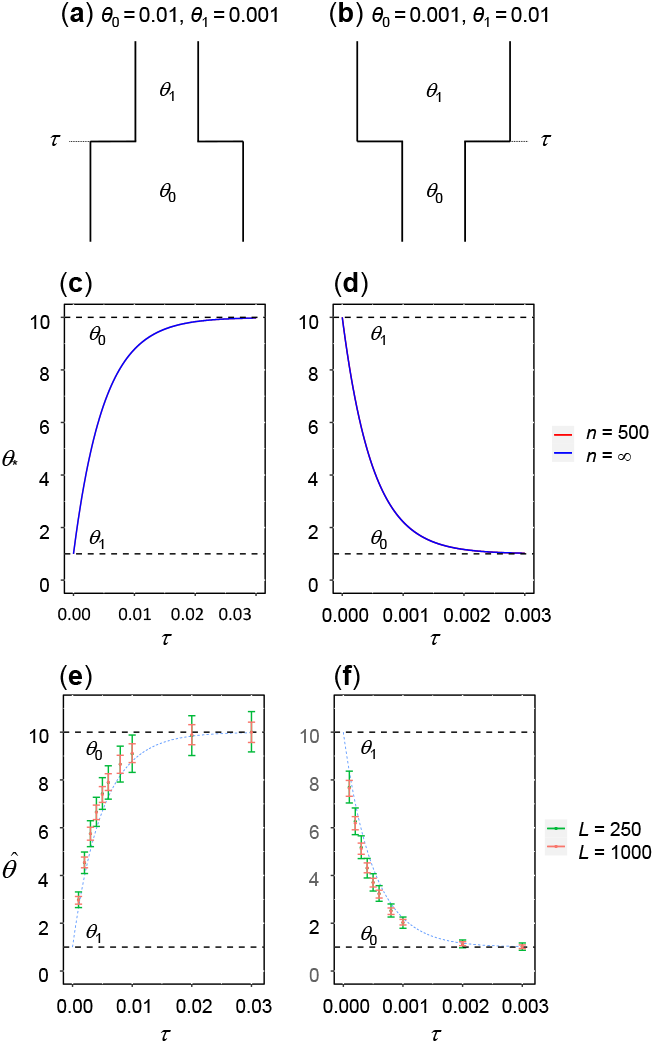
(**a, b**) Demographic models of population size change, with *θ*_0_ and *θ*_1_ over the time periods (0, *τ*) and (*τ*, ∞ ), respectively. In (**a**), *θ*_0_ = 0.01 and *θ*_1_ = 0.001 while in (**b**), *θ*_0_ = 0.001 and *θ*_1_ = 0.01. (**c, d**) Limiting estimates of *θ* under the model of constant size (*θ*_∗_ of eqs. 5 & 12, ×10^3^) when data of infinitely many loci (*L* = ∞ ), each of two sequences with *n* = ∞ or 500 sites, are generated under the two-*θ* models of **a** and **b**. Estimates for *n* = ∞ and 500 are indistinguishable. (**e, f**) Average posterior means and the 95% HPD CIs for *θ* from BPP analysis of data of *L* = 250 or 1000 loci simulated under models of **a & b**. Blue dashed lines represent the asymptotic estimates from panels **c & d**.

## Materials and Methods

### Comparison of models of gene flow for a species triplet and inference of ghost introgression

To compare the models of gene flow of Figure 2**a**-**c**: ghost introgression, inflow, and outflow, we implement a Bayesian testing approach. We conduct a Markov chain Monte Carlo (MCMC) analysis under an MSC-I model that is more general and includes all the three models of Figure 2**a**–**c** as special cases. This involves three introgression probabilities, *φ*_*H*_, *φ*_*x*_, and *φ*_*y*_, for ghost introgression, inflow, and outflow, respectively (Fig. 2**d**). We then use the MCMC sample under the general model to test whether each introgression probability is different from the null value of 0. As the hypotheses being tested are nested, the Bayes factor is given by the Savage-Dickey (S-D) density ratio (Ji *et al*., 2023). In other words, the Bayes factor in support of the alternative hypothesis of gene flow (*H*_1_ : *φ >* 0) against the null hypothesis of no gene flow (*H*_0_ : *φ* = *φ*_0_ = 0) is

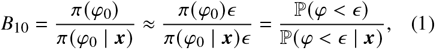

where *π* (*φ*_0_) and *π* (*φ*_0_| ***x***) are the prior and posterior probability densities of *φ* under *H*_1_, evaluated at the null value *φ*_0_ = 0 (Ji *et al*., 2023), and where the small value *ϵ* is fixed at *ϵ* = 0.01*σ* = 0.00289, with 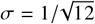 to be the standard deviation of the uniform prior *φ* ∼ *U* (0, 1).

In practice the prior probability 𝕡 (*φ < ϵ*) = *ϵ* while the posterior probability 𝕡 (*φ < ϵ*| ***x***) is estimated by the proportion of MCMC samples in which *φ < ϵ* . The S-D approach requires only one run of the MCMC algorithm under the model of Figure 2**d** and is a few hundred times more efficient computationally than calculation of marginal likelihood values under the three models using thermodynamic integration combined with Gaussian quadrature, implemented in BPP (Gelman and Meng 1998; Lartillot and Philippe, 2006; Rannala and Yang 2017) and used by Pang and Zhang (2024).

Following Pang and Zhang (2024), we simulated datasets under each of the three models of Figure 2**a-c**. The values of parameters used were *τ*_*R*_ = 2*θ, τ*_*S*_ =*θ, τ*_*H*_ = *τ*_*y*_ = 0.5*θ*, with *θ* = 0.01, while in **a**, *τ*_0_= 3*θ*. Note that time is scaled by mutations so that both *τ* and *θ* are measured in units of expected number mutations per site, and one coalescent time unit (i.e.. 2*N* generations for a diploid species of size *N*) is *θ*/2. The introgression probability used was *φ* = 0.2 in **a-c**. There were two data sizes, with *L* = 250 or 1000 loci, with *S* = 4 sequences per species per locus, and with the sequence length to be 500 sites. The number of replicates was 100. Each dataset was generated using the `simulate` option in BPP (Yang, 2015; Flouri *et al*., 2018), which simulates the gene tree (both tree topology and branch lengths) for each locus, and then generates the sequence alignment by evolving sequences on the tree under the JC mutation model (Jukes and Cantor, 1969).

Each dataset was analyzed under the general model of Figure 2**d** using BPP to test the introgression probabilities (*φ*_*H*_, *φ*_*x*_, and *φ*_*y*_). The JC mutation model was assumed. A gamma prior was assigned on the population size parameter, *θ* ∼ G(2, 200) with the prior mean 0.01. The same population size was assumed for a branch before and after introgression (using the `thetamodel = linked-msci` option in BPP); for example, *Sx* and *xB* on the species tree (Fig. 2**d**) were considered one branch and assigned the same *θ*. The age of the root of the species tree was assigned another gamma prior, *τ*_*0*_ ∼ G(2, 67) with mean 0.03 (Fig. 2**d**) while the other node ages have a uniform-Dirichlet distribution (Yang and Rannala, 2010, eq. 2). The shape parameter *α* = 2 is used so that the gamma prior is diffuse. Datasets simulated in this paper are all large so that the prior does not have an important effect.

We used a burn-in of 4 × 10^4^ iterations, after which we took 10^5^ samples sampling every 2 iterations. Running time for each dataset was 1.5hrs using one thread.

The simulated data were also analyzed using several summary methods including HyDe, implemented using the script `run hyde.py` from https://github.com/pblischak/HyDe (Blischak *et al*., 2018), and SNaQ, implemented in the PhyloNetworks package (Solis-Lemus and Ane, 2016), with gene trees for loci inferred using RAxML with default settings (Stamatakis, 2014). SNaQ is very similar to PhyloNet/MPL (Yu and Nakhleh, 2015). HyDe uses site-pattern counts pooled across all loci, and is calculated using a concatenated alignment. As HyDe requires at least four taxa, a distant outgroup species at the divergence time 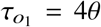 was included when data were simulated under the models of Figure 2**a**–**c**. SNaQ requires at least five taxa, so we included two distant outgroups at divergence times 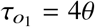, 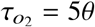. Only one sequence per species per locus was used for the summary methods.

### One species: estimation of θ for one species assuming a constant θ when θ changes over time

We develop an asymptotic theory for estimation of *θ* for one species using *S* = 2 sampled sequences (of *n* = ∞ or 500 sites) when the number of loci *L* → ∞ . The population size or *θ* is variable, being *θ*_0_ and *θ*_1_ over two time periods (0, *τ*) and (*τ*, ∞), respectively, but is assumed to be constant when the data are analyzed (Fig. 3**a&b**). The limiting value of the maximum likelihood estimate (MLE), *θ*_∗_, is analytically tractable (see the Results section for the theory). We calculated *θ*_∗_ as a function of *τ* for two scenarios: (a) *θ*_0_ = 0.01, *θ*_1_ = 0.001 and (b) *θ*_0_ = 0.001, *θ*_1_ = 0.01.

We then used simulation to examine Bayesian estimation of *θ* using more than two sequences per locus but with a finite number of loci. We used *S* = 4 sequences and *L* = 250 or 1000 loci, with the sequence length to be *n* = 500 sites. The number of replicates was 100. Each dataset was analyzed using BPP assuming a constant *θ*. The JC mutation model was assumed. A gamma prior was assigned, *θ* ∼ G(2, 200) with the prior mean 0.01 for (a) and *θ* ∼ G(2, 2000) with the mean 0.001 for (b) (Fig. 3**a&b**). We used a burn-in of 4 × 10^4^ iterations, after which we took 10^5^ samples sampling every 2 iterations. Running time for each analysis was 0.5 hrs using one thread.

### Two species: estimation of the rate of gene flow between sister species using BPP

We simulated data under the MSC-I and MSC-M models of Figure 4 **a&b**, with gene flow between two sister species, and analyzed the data under the MSC-I model (Fig. 4**a**). Species represented by thin branches had the population size *θ*_0_ = 0.001 while those for thick branches had a large size *θ*_1_ = 0.01. Note that in this study, the terms species and population are used interchangeably. Species split times were *τ*_*R*_ = 10*θ*_0_, *τ*_*S*_ = 2*θ*_0_, *τ*_*T*_ = *θ*_0_, while introgression time was *τ*_*x*_ = *τ*_*y*_ = 1.5*θ*_0_. We used eleven introgression probabilities in the MSC-I model (*φ* = 0, 0.01, 0.05, 0.1, 0.2, 0.3, 0.4, 0.5, 0.6, 0.7, 0.8) and nine migration rates in MSC-M (*M* = 0, 0.01, 0.03, 0.05, 0.1, 0.2, 0.3, 0.4, 0.5). Note that in the MSC-M model, the population migration rate, *M*_*xy*_ = *m*_*xy*_ *N*_*y*_, is the expected number of migrants from *x* to *y* per generation, with *m*_*xy*_ to be the proportion of migrants in population *y* from *x*.

**Fig. 4.**
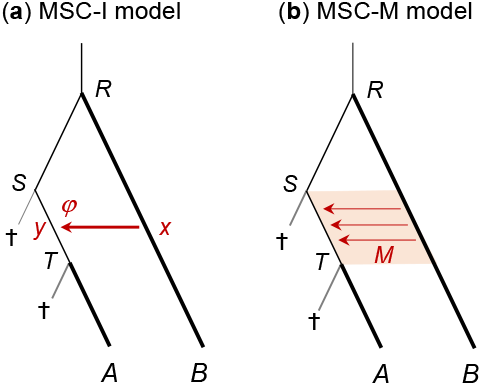
(**a**) Introgression (MSC-I) and (**b**) migration (MSC-M) models used to simulate sequence data. Branch thickness represents population size, with *θ*_0_ = 0.001 and *θ*_1_ = 0.01 for thin and thick branches, respectively. Nodes *S* and *T* are created via extinct ghost species to specify different *θ*s for different segments of the branch *R A*, but no sequences were generated for the ghost species.

We generated 100 replicate datasets. Each consisted of *L* = 250 or 1000 loci, with *S* = 4 sequences per species per locus, and with the sequence length to be *n* = 500 sites.

Each replicate dataset was analyzed under the MSC-I model (Fig. 4**a**) using BPP, with the correct donor and recipient populations identified. The root age was assigned the gamma prior *τ*_*R*_ ∼ G(2, 200) with prior mean 0.01, and population sizes were assigned the prior *θ* ∼ G(2, 2000) with mean 0.001. The same population size parameter was assumed for a branch before and after introgression. The introgression probability was assigned the uniform prior, *φ* ∼ beta(1, 1). We used a burn-in of 4 × 10^4^ iterations, after which we took 10^5^∼ samples sampling every 2 iterations. Running time was ~ 1 hour using four threads.

As a reference for comparison, we found it useful to simulate another set of data assuming the same population size for all species (*θ* = 0.001), and analyze the data assuming one population size for the whole tree (using the `thetamodel = linked-all` option in BPP). The other settings were the same as above for simulation and analysis of data assuming different *θ*s. With one *θ* for all, the issue of incomplete taxon sampling is irrelevant, and the analysis model is correctly specified.

### Four species: estimation of the rate of gene flow using BPP and summary methods

The MSC-I and MSC-M models for four species of Figure 5**a&b** were used to simulate data, which were analyzed using both BPP and summary methods. As in the case of two species, thin and thick branches on the tree had different population sizes (*θ*_0_ = 0.001 and *θ*_1_ = 0.01, respectively), and an unsampled ghost species was used to break the branch *T B* into two segments with different population sizes. Species split times were *τ*_*R*_ = 4*θ*_0_, *τ*_*S*_ = 3*θ*_0_, *τ*_*T*_ = 2*θ*_0_, *τ*_*U*_ = *θ*_0_, while introgression time was *τ*_*x*_ = *τ*_*y*_ = 1.5*θ*_0_. As in the case of two species, each dataset consisted of *L* = 250 or 1000 loci, with *S* = 4 sequences per species per locus, and with the sequence length to be *n* = 500 sites. The number of replicates was 100.

**Fig. 5.**
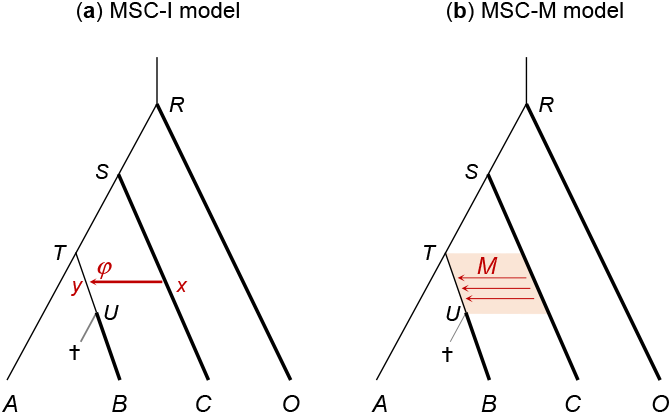
(**a**) Introgression (MSC-I) and (**b**) migration (MSC-M) models used to simulate data to compare bpp and summary methods for estimating the rate of gene flow.

**Fig. 6.**
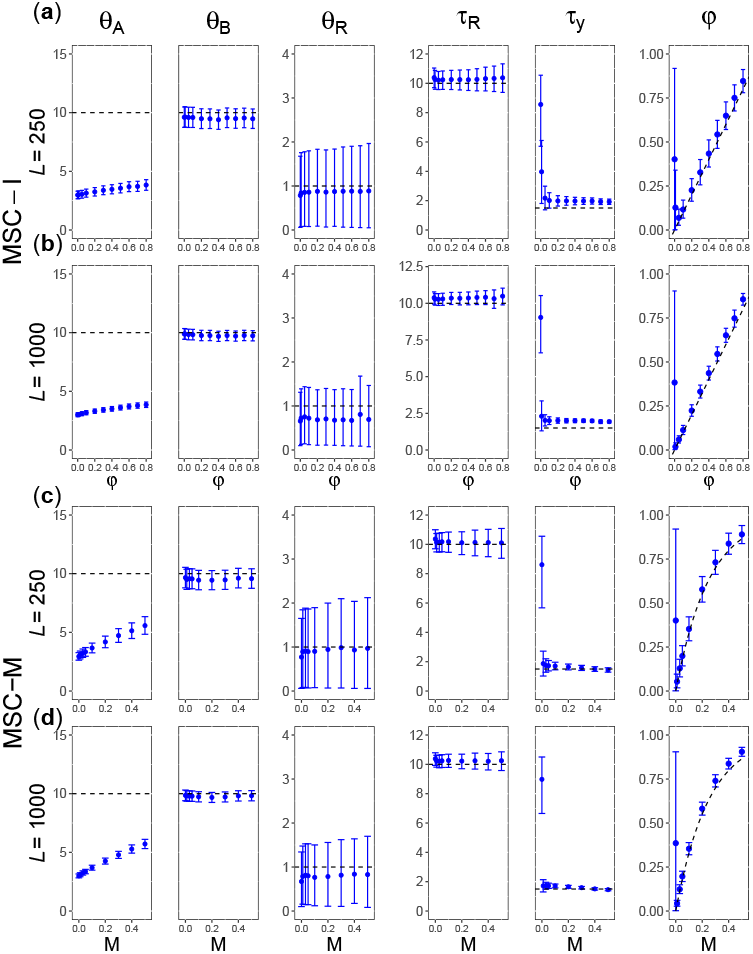
[2s-diff-diff] (**a, b**) Average posterior means and the 95% HPD CIs of parameters when the data are simulated and analyzed under the MSC-I model (Fig. 4**a**) assuming different *θ*s for species, plotted against the introgression probability (*φ*). (**c, d**) Parameter estimates under MSC-I when data are simulated under MSC-M (Fig. 4**b**), plotted against the migration rate (*M*). Estimates of *θ* and *τ* are mutipled by 1000. Black dashed lines represent the true value in **a&b** or expected value (*φ*_0_) in **c&d**.

Each replicate dataset was analyzed under the MSC-I model of Figure 5**a**. Different species on the phylogeny were assigned independent population sizes (*θ*), although the same branch before and after introgression was assigned the same size. The settings for running BPP were the same as in the analysis of the two-species data. The priors were *θ* ∼ G(2, 400), and *τ*_*R*_ ∼ G(2, 500), and *φ* ∼ beta(1, 1). We used a burn-in of 4 × 10^4^ iterations, after which we took 10^5^ samples sampling every 2 iterations.

To help interpret the simulation results, we simulated another set of data assuming one population size for all species (*θ* = *θ*_0_ = 0.001), and analyze the data assuming one *θ* for all species. All other settings were the same as above.

The simulated data were also analyzed using HyDe and SNaQ (equivalent to PhyloNet/MPL) to infer gene flow and to estimate the introgression probability. Both methods produce point estimates of *φ* only, while BPP provides in addition a measure of uncertainty in posterior CIs. Note that those summary methods were developed under the assumption of one population size (*θ*) for all species on the phylogeny.

### Six species: inference of gene flow on a species phylogeny using BPP and f -branch

We simulated sequence data under the MSC-I model for six species of Figure 1**a** and analyzed the data using BPP and the *f* -branch method based on species triplets. We used *θ*_0_ = 0.001 for thin branches and *θ*_1_ = 0.01 for thick branches on the phylogeny. Species split times were *τ*_*R*_ = 5*θ*_0_, *τ*_*S*_ = 4*θ*_0_, *τ*_*T*_ = 3*θ*_0_, *τ*_*U*_ = 2*θ*_0_, *τ*_*V*_ = *θ*_0_, 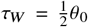, while introgression times were *τ*_*x*_ = *τ*_*y*_ = 2.5*θ*_0_, *τ*_*u*_ = *τ*_*v*_ = 1.5*θ*_0_, and *τ*_*z*_ = *τ*_*w*_ = 0.5*θ*_0_ . The introgression probabilities were *φ*_*y*_ = 0.3, *φ*_*u*_ = 0.2 and *φ*_*w*_ = 0.1. As in the case of two or four species, we used *L* = 250 or 1000 loci, with *S* = 4 sequences per species per locus, and with the sequence length to be *n* = 500. The number of replicates was 100.

We used BPP to analyse the data under the MSC-I model (Fig. 1**a**). The same settings were used as above. The priors were *θ* ∼ G(2, 400), and *τ*_*R*_ ∼ G(2, 400), and *φ* ∼ beta(1, 1). We used a burn-in of 4 × 10^4^ iterations, after which we took 10^5^ samples sampling every 2 iterations. Running time for analyzing each dataset was ∼1.5hrs using 4 threads.

We also conducted the Bayesian test of gene flow, using the MCMC sample collected under the MSC-I model to calculate the Bayes factor via the S-D density ratio (Ji *et al*., 2023).

We also simulated a set of data assuming one population size for all species (*θ* = 0.001), and analyzed the data using BPP assuming one *θ* for all species. The other settings were the same as in the analysis of data simulated under different *θ*s.

The simulated data were also analyzed using the *f* -branch approach, implemented in the Dsuite software (Malinsky *et al*., 2021). The correct species tree of Figure 1**a** was provided. The approach first analyzes all triplets, using a triplet method such as *f*_4_ (Patterson *et al*., 2012) and HyDe (Blischak *et al*., 2018), and then assembles the triplet estimates by attempting to move introgression events onto ancestral branches (Malinsky *et al*., 2018, 2021).

## Results

### Comparison of models of gene flow for a species triplet and inference of ghost introgression

We simulated datasets under the three models of Figure 2**a**–**c**, ghost introgression, inflow, and outflow, and fitted the general model of Figure 2**d** to test the presence of introgression events, with the Bayes factor calculated using the S-D density ratio (eq. 1). The results are summarized in table 1.

**Table 1.** Power of Bayesian and summary methods to detect introgression, measured as percentages of replicate datasets in which the test supports each of the three introgression events in the general model of Figure 2**d** in datasets simulated under the three models of Figure 2**a**–**c**

| True model | (a) Ghost |  |  | (b) Inflow |  |  | (c) Outflow |  |  |
| --- | --- | --- | --- | --- | --- | --- | --- | --- | --- |
| | $\varphi_H$<br>(ghost) | $\varphi_x$<br>(inflow) | $\varphi_y$<br>(outflow) | $\varphi_H$<br>(ghost) | $\varphi_x$<br>(inflow) | $\varphi_y$<br>(outflow) | $\varphi_H$<br>(ghost) | $\varphi_x$<br>(inflow) | $\varphi_y$<br>(outflow) |
| ***L = 250 loci*** |  |  |  |  |  |  |  |  |  |
| $0 < B_{10} < 0.0101$ (very strong rejection) | 0 | **97** | **100** | **33** | 0 | **90** | **41** | **77** | 0 |
| $0.0101 < B_{10} < 0.0526$ (strong rejection) | 0 | **3** | **0** | **55** | 0 | **6** | **51** | **20** | 0 |
| $0.0526 < B_{10} < 1$ (rejection) | 0 | 0 | 0 | 11 | 0 | 4 | 6 | 3 | 0 |
| $1 < B_{10} < 19$ (support) | 0 | 0 | 0 | 1 | 0 | 0 | 2 | 0 | 0 |
| $19 < B_{10} < 99$ (strong support) | 0 | 0 | 0 | 0 | 0 | 0 | 0 | 0 | 0 |
| $99 < B_{10} < \infty$ (very strong support) | **100** | 0 | 0 | 0 | **100** | 0 | 0 | 0 | **100** |
| *Summary methods* |  |  |  |  |  |  |  |  |  |
| HyDE | 0 | 100 | 0 | 0 | **100** | 0 | 0 | 100 | 0 |
| SNAQ and PHYLONET/MPL | **95** | 5 | 0 | 30 | **28** | 41 | 17 | 11 | **72** |
| ***L = 1000 loci*** |  |  |  |  |  |  |  |  |  |
| $0 < B_{10} < 0.0101$ (very strong rejection) | 0 | **100** | **100** | **84** | 0 | **100** | **73** | **97** | 0 |
| $0.0101 < B_{10} < 0.0526$ (strong rejection) | 0 | 0 | 0 | **22** | 0 | 0 | **13** | **3** | 0 |
| $0.0526 < B_{10} < 1$ (rejection) | 0 | 0 | 0 | 4 | 0 | 0 | 4 | 0 | 0 |
| $1 < B_{10} < 19$ (support) | 0 | 0 | 0 | 0 | 0 | 0 | 1 | 0 | 0 |
| $19 < B_{10} < 99$ (strong support) | 0 | 0 | 0 | 0 | 0 | 0 | 0 | 0 | 0 |
| $99 < B_{10} < \infty$ (very strong support) | **100** | 0 | 0 | 0 | **100** | 0 | 0 | 0 | **100** |
| *Summary methods* |  |  |  |  |  |  |  |  |  |
| HyDE | 0 | 100 | 0 | 0 | **100** | 0 | 0 | 100 | 0 |
| SNAQ and PHYLONET/MPL | **100** | 0 | 0 | 15 | **50** | 35 | 6 | 8 | **86** |

Note.— Bayesian test is based on the Bayes factor, $B_{10}$ , calculated using the S-D density ratio with $\epsilon_\phi = 0.00289$ . The cutoffs, $B_{10} = 0.0101, 0.0526, 1, 19$ , and $99$ , correspond to the posterior probability for the model of gene flow $\mathbb{P}(H_1|x) = 0.01, 0.05, 0.5, 0.95$ , and $0.99$ , respectively. Strong support for the true model and strong rejection of the wrong model (i.e., non-existent gene-flow events) are highlighted in bold. Each column sums to 1. HyDE assumes inflow only, and the support for inflow is significant when the $p$ -value is $< 5\%$ . SNAQ (equivalent to PHYLONET/MPL in the case of four species) ranks the three introgression models by the pseudo-likelihood based on the counts of gene tree topologies.

We found strong support for the correct introgression events and often strong rejection of nonexistent introgression events (table 1). For example, even in small datasets of *L* = 250 loci, simulated under the model of ghost introgression, the correct model of ghost introgression (*φ*_*H*_) was very strongly supported (with *B*_10_ *>* 99) in 100% of datasets, while inflow (*φ*_*x*_) was very strongly rejected in 97% of datasets (with *B*_10_ *<* 0.0101), and outflow (*φ*_*y*_) was very strongly rejected in 100% of them. Note that strong rejection of the alternative hypothesis is possible in the Bayesian test.

Similarly the correct model was strongly supported in 100% of datasets simulated under the inflow or outflow models, while introgression in the opposite direction (as well as introgression from the ghost species) was strongly rejected. The result is consistent with the previous finding that use of the bidirectional introgression model allows the Bayesian approach to identify the direction of gene flow (Thawornwattana *et al*., 2023).

The simulated datasets were also analyzed using the summary methods HyDe and SNaQ. HyDe assumes inflow only, and provides significant support for inflow when the *p*-value is *<* 5%. In every dataset simulated here, HyDe found significant evidence for inflow, irrespective of the true model.

SNaQ is equivalent to PhyloNet/MPL in the case of four species, and ranks the three introgression models by the pseudo-likelihood based on the counts of gene tree topologies. It does not provide significance values, so the results are not directly comparable with those of BPP. SNaQ selected the correct model in 95%, 28%, and 72% of datasets, when the true model was ghost introgression, inflow, and outflow, respectively. However, the results were noted to depend on the divergence times for the two outgroups, and at 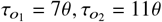, the corresponding proportions were 32%, 48%, and 17%. As the models under comparison are unidentifiable, the result may depend on the idiosyncrasies of software implementation. The results are consistent with Pang and Zhang (2024, Fig. 3).

### One species: estimation of θ for one species assuming a constant θ when θ changes over time

To examine the impact of changing population sizes due to taxon exclusion, we consider first the simple problem of estimating the population size parameter (*θ*) for a single population, when *θ* changes over time, being *θ*_0_ and *θ*_1_ over two time periods (0, *τ*) and (*τ*, ∞), respectively (Fig. 3**a**). We deal with the case of two sequences per locus first, which is analytically tractable.

Let *θ* (*t*) be the population size time *t* ago. The density for the coalescent time (*t*) between two sequences is then

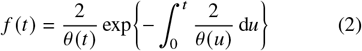

(Slatkin and Hudson, 1991; Griffiths and Tavaré, 1994). With the two-*θ* model of Figure 3**a**, this simplifies to

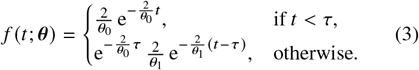

Suppose coalescent times at *L* loci, *t*_*i*_, *i* = 1,, · · · *L*, are available and used to estimate *θ* under the model of a constant population size (*H*_0_). The coalescent time has the density

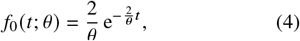

giving the likelihood 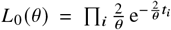 .The MLE under *H*_0_ is simply 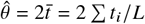.

When *L* → ∞, the MLE *θ* approaches the limiting value *θ*_∗_, which minimizes the Kullback–Leibler (KL) divergence

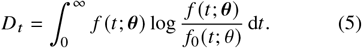

Note that *θ*_∗_ is also the limiting value of the Bayesian estimate (posterior mean) as *L* → ∞ . It is known as the *best-fitting parameter value* or *pseudo-true parameter value* under *H*_0_. In eq. 5, *f* (*t*; ***θ***) represents the data while *f*_0_(*t*; *θ* ) represents the fitting model, and minimization of equation 5 is equivalent to maximization of the average log likelihood per datum under the fitting model *H*_0_,

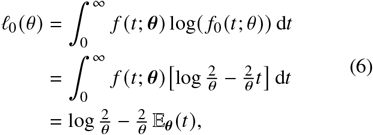

Giving

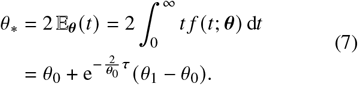

This is a weighted average of *θ*_0_ and *θ*_1_, and the weight 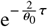 for *θ* is the probability that the coalescence occurs over (*τ*, ∞ ) when the population size is *θ*_1_ (Fig. 3).

Treating coalescent times as data is equivalent to assuming infinitely long sequences. Next we consider finite sequences with *n* sites. We assume the infinite-sites mutation model. The number of mutations (*x*) between two sequences of length *n*, given the coalescent time *t*, has the Poisson distribution

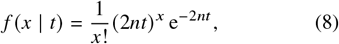

where 2*nt* is the expected number of mutations between the two sequences. The unconditional probability is an average over the coalescent time,

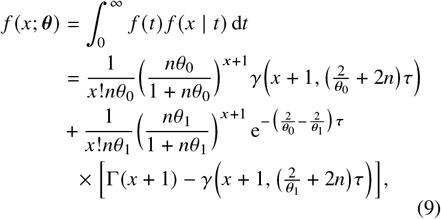

Where

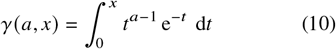

is the incomplete gamma function and Γ *a* = *γ* (*a*, ∞) is the gamma function, with Γ (*x* + 1) = *x*! as *x* is a non-negative integer.

Under *H*_0_ with a constant *θ, x* has the density

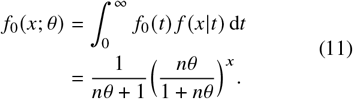

In other words, the number of segregating sites (*x*) has the geometric distribution (Watterson, 1975).

Again by minimizing the K-L divergence,

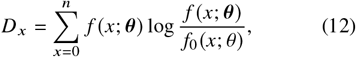

we obtain the best-fitting parameter value *θ*_∗_ under *H*_0_, as a function of (*τ, θ*_0_, *θ*_1_) in the generating model and the sequence length (*n*). In this case the solution is not analytical. Instead we use the BFGS algorithm implemented in paml (Yang, 2007) to find *θ*_∗_ to minimize *D* _*x*_ (eq. 12), with Gaussian quadarature used to calculate the 1-D integral numerically, using 64 quadrature points.

The limiting estimate *θ*_∗_ is plotted against *τ* in Figure 3**c** &**d** for two sets of population sizes: (a) *θ*_0_ = 0.01, *θ*_1_ = 0.001 and (b) *θ*_0_ = 0.001, *θ*_1_ = 0.01. The results for *n* = ∞ and *n* = 500 are indistinguishable. In the special case of *τ* = 0, we have *θ*_∗_ = *θ*_1_ while if *τ* = ∞, *θ*_∗_ = *θ*_0_. When 0 *< τ <, θ*_∗_ is an average between *θ*_0_ and *θ*_1_, with weights to be the probabilities that the coalescent occurs in the two time periods (eq. 7). At 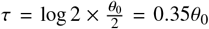, half of the coalescent events occur before reaching *τ* (that is, with *t < τ*), so that 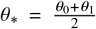 of coalescent events occur before *τ*, so that *θ*_∗_ is dominated by *θ*_0_.

We then simulated finite datasets of *L* = 250 or 1000 loci, with posterior means and the 95% highest posterior density (HPD) credibility intervals (CIs) shown in Figure 3**e&f**. There was more uncertainty in estimates for *L* = 250 than for *L* = 1000, but the means were similar between the two data sizes, which may be expected to be close to the limiting value *θ*_∗_ when *L* → ∞ . While the mean value is a weighted average of *θ*_0_ and *θ*_1_, it is closer to *θ*_0_ than in the case of *S* = 2 sequences. This is because with more sequences in the sample, a greater proportion of coalescent events occurs in the recent time interval (0, *τ*), so that the estimate *θ*_∗_ will be closer to *θ*_0_. Overall, the results for data of four sequences (Figs. 3**e&f**) were similar to those for two sequences (with either *n* = ∞ or *n* = 500 sites) (Figs. 3**c&d**).

### Two species: estimation of the rate of gene flow between sister species

We simulated data under the MSC-I and MSC-M models for two species of Figure 4 and analyzed them under the MSC-I model using BPP. Summary methods considered in this study cannot infer gene flow between sister species and were not used.

### One population size for all species

First we describe the simulation assuming one population size for all species (Fig. S1). For data simulated under the MSC-I model, the correct model is specified and incomplete taxon sampling has no effects except for affecting the information content (Fig. S1**a&b**). The results represent the best-case scenario and thus provide a reference for comparison. With the same *θ* assumed for all populations on the species tree, *θ* was very accurately and precisely estimated, even at *L* = 250 loci. Other parameters (*τ*_*R*_, *τ*_*y*_, *φ*) were well estimated as well when the rate of gene flow was not very low. However, when the true rate was very low (*φ* = 0 or 0.01), the introgression probability (*φ*) had large uncertainties, with the CI covering almost the whole range (0, 1) for the parameter. The other parameters were affected as well.

For data simulated under MSC-M, the mode of gene flow is misspecified but the assumptions about population sizes are all satisfied (Fig. S1**c&d**). The results were strikingly similar to those of Figure S1**a&b**, where the MSC-I model was used both to simulate and to analyze data. The misspecification of the mode of gene flow had very little impact. Parameter *θ* was very accurately estimated. The other parameters were well estimated as well, except when gene flow was absent (*M* = 0). When *M* = 0, the species split time, the introgression time, and the introgression probability involved large uncertainties (Fig. S1**c&d**).

The poor estimates at very low rates of gene flow were noted by Huang *et al*. (2022, Fig. 3, *M*), where data were simulated under the MSC-M model with migration between sister species and analyzed under the MSC-I model. Here the same pattern is seen whether the MSC-I or MSC-M model is used to generate data.

When there is continuous migration from populations *x* to *y* over time period Δ*τ*, the total amount of gene flow expected under the MSC-M model may be measured by the probability that a sequence from the recipient population *y* is traced back to the hybridizing population *x*, given as

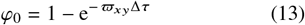

where *ϖ*_*xy*_ = *m*_*xy*_ /*μ* = 4*M*_*xy*_ /*θ*_*y*_ is the mutation-scaled migration rate (Huang *et al*., 2022, eq. 10).

When the migration rate *M* increases, the expected amount of gene flow *φ*_0_ increases, and the estimated *φ* under the MSC-I model closely tracked the expectation *φ*_0_, suggesting that the MSC-I model was able to recover nearly all the gene flow that occurred in the data (with the exception at the low rates with large uncertainty in 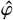) (Fig. S1**c&d**).

### Different population sizes with incomplete taxon sampling

Results for data simulated assuming different population sizes for species on the phylogeny (Fig. 4) are summarized in Figure 6**a&b** for data simulated under the MSC-I model and in Figure 6**c&d** for data simulated under MSC-M. In the simulation models (Fig. 4**a&b**), the recipient population (branch *R A*) is represented by three segments with different population sizes, a small size *θ*_0_ = 0.001 for *RS* and *ST* and a large size *θ*_1_ = 0.01 for *T A*. When the data were analyzed, one population size was assigned to the whole branch *R A* (or *RST A*).

As in the analysis of data assuming one *θ* for all species, the results were very similar between data simulated under MSC-I and under MSC-M (Fig. 6**a**-**d**), and the misspecification of the mode of gene flow had little impact. Our analysis of the case of one species (Fig. 3**a-f**) suggests that here the estimated *θ* _*A*_ may be an average of *θ*_0_ = 0.001 and *θ*_1_ = 0.01 in the true model, with weights corresponding to the proportions of coalescent events that occur in populations *RT* (with *θ*_0_ = 0.001) and *T A* (*θ*_1_ = 0.01). The increase of 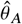 with the increase of the rate of gene flow (*φ* in MSC-I and *M* in MSC-M) may be similarly explained, as a high *B* → *A* rate of gene flow means few *A* lineages enter and coalesce in population *RSY* . Population size *θ*_*B*_ is unaffected by taxon sampling or gene flow and was well estimated. The species split time and the population size at the root (*τ*_*R*_, *θ*_*R*_) are poorly estimated, compared with the case of one *θ* for all populations (Fig. S1).

The introgression probability (*φ*) was well estimated except at very low rates of gene flow (*φ* = 0 and 0.01 in MSC-I and *M* = 0 in MSC-M, Fig. 6). When the data were generated under MSC-M and analyzed under MSC-I, the estimate 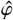 was close to the expectation *φ*_0_ (Fig. 6**c&d**), suggesting that the MSC-I model recovered almost all gene flow that occurred in the data. The estimates were nearly as good as those for data simulated assuming the same *θ* (Fig. S1). This was the case even when the population size for the recipient species (*θ* _*A*_) was incorrect by a few folds (note that the true size is *θ*_0_ = 0.001 for *RST* and *θ*_1_ = 0.01 for *T A*). The results confirm our expectation that genomic data may be informative about the proportion of immigrants rather than the number of immigrants (see Introduction).

### Four species: estimation of the rate of gene flow by bpp and quartet summary methods

We simulated data under the MSC-I and MSC-M models for four species of Figure 5 and analyzed the data using BPP under the MSC-I model, and also using summary quartet methods. Again we discuss the results for data simulated assuming one population size for all species first.

#### One population size for all species

The results for BPP are summarized in Figure S2. In Figure S2**a&b**, the MSC-I model was used both to simulate and analyze data, with no model misspecification. The results thus represent the best-case scenario. The posterior means for species split times and introgression times (*τ*_*R*_, *τ*_*S*_, *τ*_*T*_, *τ*_*Y*_ ), and the population size (*θ*) were all close to the true values and the CIs were narrow and included the true values. When the true *φ* was very low, estimates of the introgression time *τ*_*y*_ involved large uncertainties. This can be explained by the rarity of introgression events on the gene trees or in the sequence data: if an event is rare, it will be hard to estimate the time of the event. In contrast, at high *φ*, the split time *τ*_*T*_ involved large uncertainties. This is apparently due to the negative correlation between *τ*_*T*_ and *φ*: a high rate of gene flow with an ancient split may fit the data nearly equally well as a low rate of gene flow and recent split. Overall, the introgression probability (*φ*) was well estimated. Asymptotic theory predicts that quadrupling the amount of data reduces the CI by a half, a prediction that roughly holds when we compare the CIs for *L* = 250 and 1000.

In Figure S2**c&d**, data were simulated under the MSC-M or secondary-contact model of Figure 5**b** and analyzed under the MSC-I model of Figure 5**a**, so that the mode of gene flow was misspecified. Species split times and the population size were well estimated, as in Figure S2**a&b**. The estimated introgression time (*τ*_*y*_) was less than the average migration time, that is, 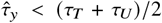. This is because recent migration events on the gene trees close to *τ*_*U*_ in the data can only be explained by a recent introgression time in the MSC-I model (Huang *et al*., 2022).

The amount of gene flow predicted in the MSC-M model (eq. 13) gave a close match to estimates of *φ* in MSC-I from BPP (Fig. S2**c&d**), suggesting that the MSC-I model recovered almost all of the gene flow that occurred in the data simulated under the MSC-M model despite the misspecification of the mode of gene flow.

The simulated data were also analyzed using the summary methods HyDe and SNaQ to estimate the introgression probability (Fig. 7). All estimates were well-behaved, although HyDe appeared to overestimate *φ* slightly. The bias may be due to the fact that HyDe assumes a hybrid-speciation model (that is, a symmetrical inflow model with *τ*_*T*_ = *τ*_*y*_ = *τ*_*x*_ in Figure 5**a**) whereas in the true model *τ*_*T*_ *> τ*_*x*_ (Ji *et al*., 2023). Note that HyDe and SNaQ do not provide estimates for species split times or introgression time.

### Different population sizes for species on the phylogeny

Quartet data generated under different population sizes (Fig. 5**a&b**) were analyzed using BPP in two ways, assuming either different population sizes (Fig. 8) or the same population size (Fig. S3).

We first consider BPP analysis assuming different *θ*s (Fig. 8**a&b**). Note that the presence of a ghost species in the model tree (Fig. 5) means that the recipient population of gene flow is represented by branch *TUB* consisting of segments with different population sizes, so that the model is misspecified even if different *θ*s are assumed for different species in the model. As there are now far more parameters in the model, the estimates, especially population sizes for ancestral species (Fig. 8**a&b**), had much larger uncertainties than in data simulated assuming one *θ* (Fig. S2). Nevertheless, the *θ* and *τ* parameters were well estimated. Species split times (*τ*_*R*_, *τ*_*S*_, *τ*_*T*_ ) appeared to be nearly as well-estimated as in the case of one *θ* for all populations. Similar to Figure S2, *τ*_*T*_ had larger CIs at high rate of gene flow and *τ*_*y*_ had larger CIs at low rate of gene flow.

Because branches *TU* and *UB*, which had different population sizes in the true model (*θ*_0_ = 0.001 for *TU* and *θ*_1_ = 0.01 for *UB*, Fig. 5**a**), are merged into one branch and assigned one population size (*θ*_*B*_) in the analysis model, the estimated *θ*_*B*_ is expected to be a weighted average of *θ*_*B*_ = 0.01 and *θ*_*U*_ = 0.001, depending on whether sequences from *B* coalesce with each other before or after *τ*_*U*_ (Fig. 8**a&b**). Despite the poor estimation of *θ*_*B*_, estimates of *φ* were close to the true values (Fig. 8**a&b**), although they involved greater uncertainties than in the case of one *θ* for all populations (Fig. S2**a&b**). The result confirmed our expectation that it is the probability *φ*_*CB*_ that a *B* sequence is traced to population *C*, rather than the expected number of immigrants (*θ*_*B*_*φ*_*CB*_), that is well estimated from sequence data.

When the data were generated under MSC-M but analyzed under MSC-I (Fig. 8**c&d**), the results were similar to those obtained from the data simulated under MSC-I (Fig. 8**a&b**). At higher migration rate (*M* = *M*_*CB*_), the estimated *θ*_*B*_ was larger. This is apparently because *θ*_*B*_ is estimated based on coalescent events of *B* sequences along the branch *BUT*, and a high migration rate reduces the proportion of coalescent events along branch *UT* (which had the small population size, Fig. 5**b**). The estimates of *φ* were close to the expected *φ*_0_ (eq. 13), suggesting that the amount of gene flow expected under the MSC-M model was nearly correctly recovered by the MSC-I model.

We also analyzed the data of Figure 8 simulated with different *θ*s using BPP under the MSC-I model assuming one *θ* for all populations (Fig. S3). The analysis model was wrong, and the parameter estimates had large biases. While there was a positive correlation between the true introgression probability in the MSC-I model and the Bayesian estimates, the match was poor (Fig. S3**a&b**, *φ*). Estimates of introgression time were affected as well (Fig. S3**a&b**, *τ*_*y*_). When the data were simulated under MSC-M with different *θ*s but analyzed under MSC-I assuming the same *θ*, parameter estimates appeared to be less biased, with the estimated introgression probability 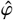 being lower than but tracking the expected value *φ*_0_ (Fig. S3**c&d**).

Overall, BPP analysis was affected far more by the assumptions of equal population size on the species phylogeny than by the mode of gene flow (MSC-I versus MSC-M).

The simulated data of Figure 8 were also analyzed using the summary methods HyDe and SNaQ (Fig. 9). Note that HyDe and SNaQ were developed under the assumption of one population size for all species on the triplet tree. The estimates of introgression probability produced by HyDe and SNaQ were poor (Fig. 9), somewhat similar to BPP estimates under the assumption of one *θ* for all populations and more different from BPP estimates under the assumption of different *θ*s. The results are in sharp contrast to the accurate estimates of Figure 7 for data simulated assuming one population size.

**Fig. 7.**
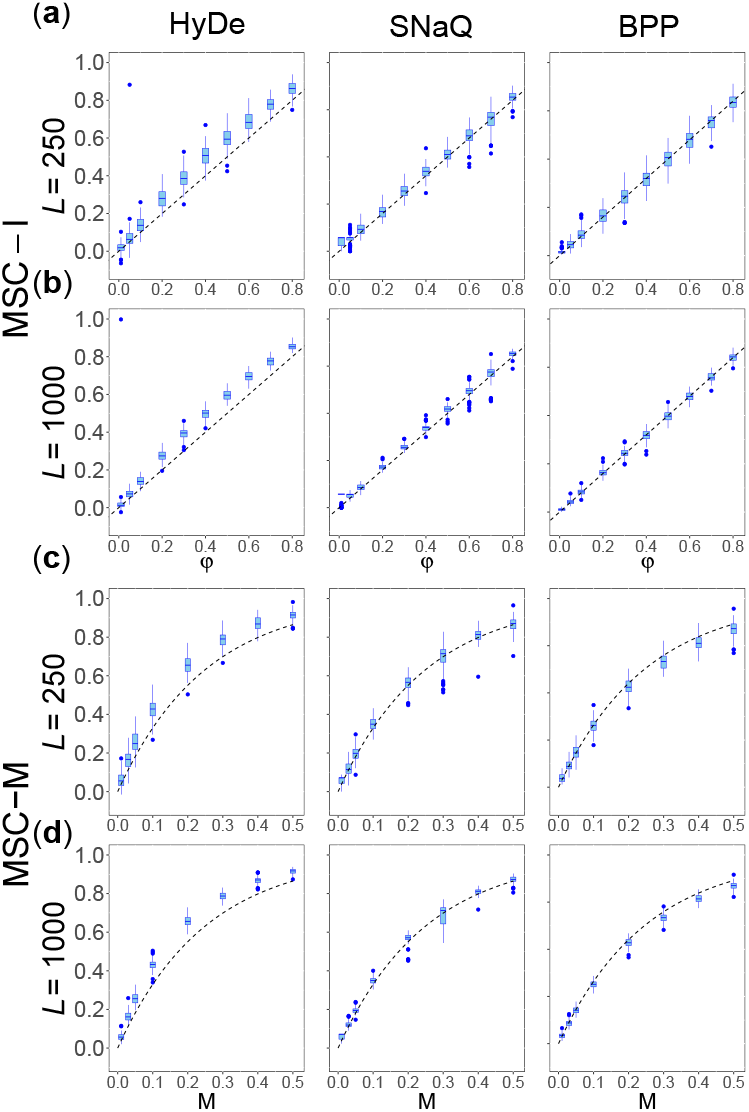
[4s-same-*θ*] Boxplots of estimates of the introgression probability *φ* using (**a**) Hyde and (**b**) SNaQ in analyses of the data of Figure S2. (**c**) Results for BPP of Figure S2 are plotted here for comparison. In a boxplot, the bar represents the median, the box the interquartile range (IQR), while the whiskers represent the minimum and maximum defined as the lower and upper quartile plus 1.5 × IQR. Note that the average 95% CI for *φ* was shown for BPP in Figure S2 whereas here the IQR based on point estimates is shown.

**Fig. 8.**
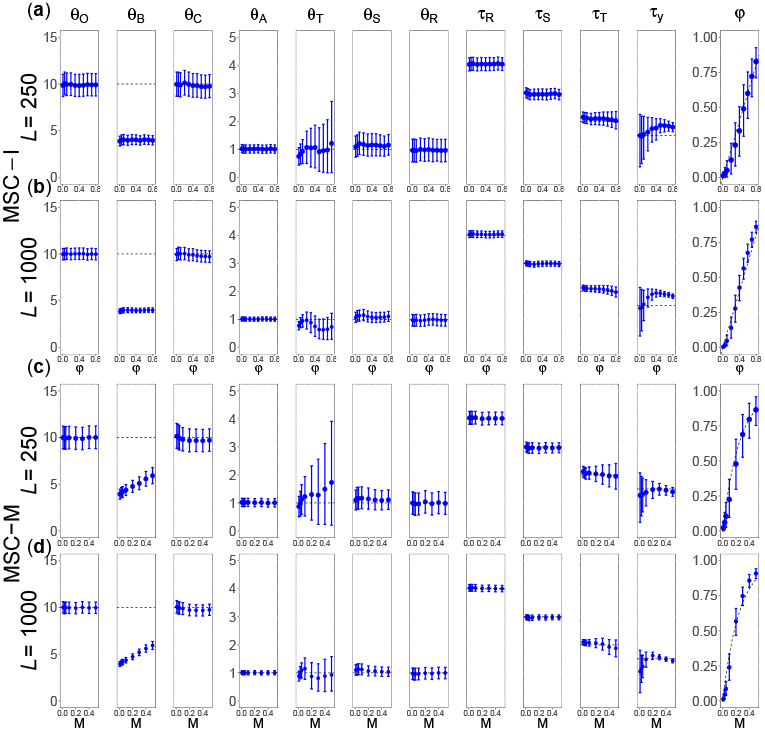
[4s-diff-diff] (**a, b**) Average posterior means and the 95% HPD CIs for parameters when data are simulated and analyzed under the MSC-I model of Figure 5a assuming different *θ*s for species. (**c, d**) Estimates under MSC-I when data are simulated under MSC-M (Fig. 5b). See legend to Figure S2.

**Fig. 9.**
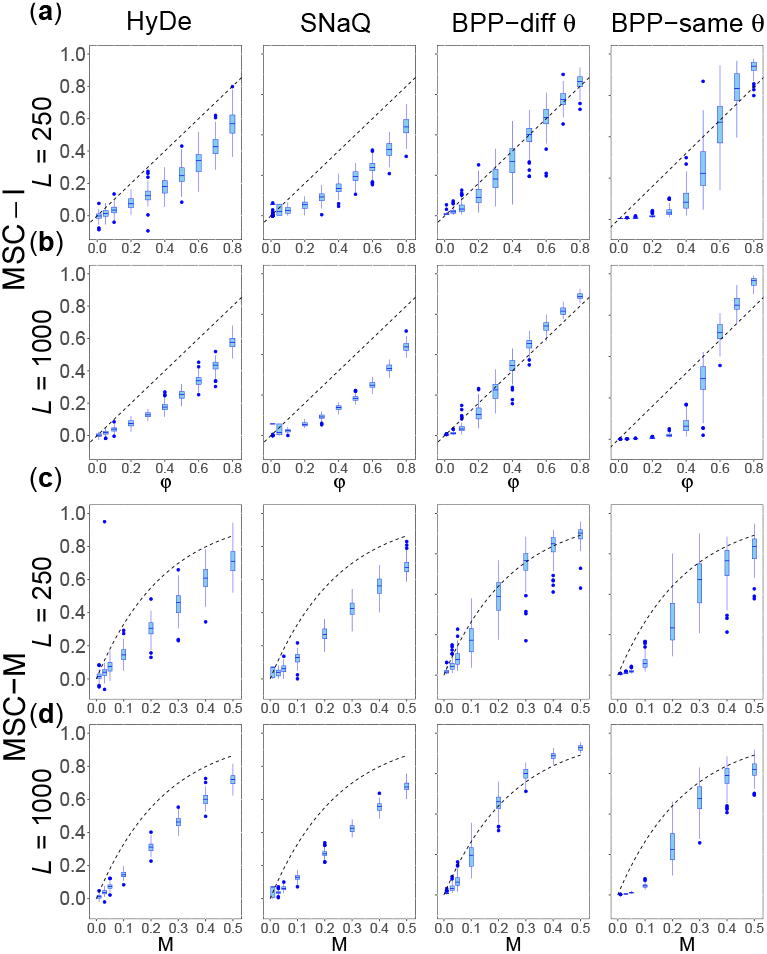
[4s-diff] Boxplots of estimates of *φ* obtained in Hyde, SNaQ, and BPP analyses of the data of Figure 8. See legend to Figure 7.

### Six species: inference of gene flow using f -branch and BPP

We simulated sequence data under the MSC-I model for six species of Figure 1**a** and analyzed the data using the *f* -branch method, implemented in Dsuite (Malinsky *et al*., 2021), to see whether it could recover the geneflow events in the true model. Two sets of data were simulated assuming either one *θ* for all populations or different *θ*s for the thin and thick branches on the species tree (Fig. 1**a**). The correct binary species tree (Fig. 1**a**) was assumed, and species *O* was used as the outgroup in all triplet analyses.

#### The f -branch approach to inferring gene flow

Here we provide an overview of the *f* -branch approach, which consists of the following steps.

First, a triplet method (e.g., *f*_4_, Patterson *et al*., 2012 or HyDe, Blischak *et al*., 2018) is used to estimate introgression probabilities for all triplets.

Second, an attempt is made to move inferred introgression events onto ancestral branches on the species tree. Consider the species tree of Figure 10, in which *A* and *B* are clades, and *b* is an ancestral branch. Selecting a species *A*_*i*_ from clade *A* and another species *B* _*j*_ from clade *B* leads to many triplets of the form *A*_*i*_ *B* _*j*_*C*, generating many *φ* estimates. The method moves the introgression event to the ancestral branch *b* with the estimate of the introgression probability given by

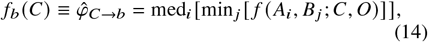

where *f A*_*i*_, *B* _*j*_; *C, O* is the estimate of *φ*_*C*→*B*_ from the *A*_*i*_ *B* _*j*_*C* triplet (using, e.g., *f*_4_), and where med stands for median and min for minimum (Malinsky *et al*., 2018).

**Fig. 10.**
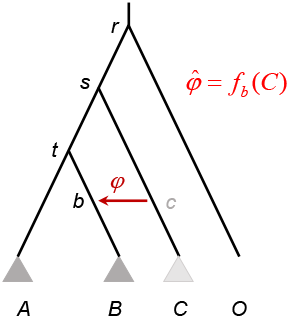
Species tree with an outgroup (*O*) to illustrate the *f* -branch approach. Selecting a species *A*_*i*_ from clade *A* and another species *B* _*j*_ from clade *B* leads to many triplets of the form *A*_*i*_ *B* _*j*_*C*, with each triplet generating an *φ* estimate. The approach then integrates the triplet estimates to produce an introgression probability into the ancestral branch 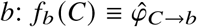.

Third, estimates generated from eq. 14 are filtered via a statistical test or an empirical cut-off, and estimates that do not pass the test are reset to 0. Dsuite uses a block-jackknife procedure to estimate the standard error of the *D*-statistic, and calculates a *Z* score or *p*-value, and uses the Holm-Bonferroni correction to control the family-wise error rate (FWER) (Malinsky *et al*., 2021). Alternatively an empirical cut-off on *φ* may be applied, for example, 0.03 (Malinsky *et al*., 2018) or 0.05 (Malinsky *et al*., 2021). If the triplet estimate does not pass the test (e.g., if the *p*-value is *>* 0.01), the *f* -branch estimate is set to 0. The rationale is that one should not claim gene flow unless the evidence is strong.

We present the *f* -branch analysis of a replicate dataset in detail in Figure 1**c**, to illustrate the procedure. The analysis produced three nonzero estimates. They were, in decreasing order, 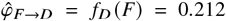 based on the *CDF* triplet, 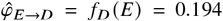 based on *CDE*, 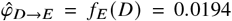 based on *FE D* (Fig. 1**c**). Among the three inferred events, the *E* → *D* gene flow clearly reflected the rates *φ*_*z*→*w*_ = 0.1 and *φ*_*v*→*u*_ = 0.2 in the true model (Fig. 1**a**). However the estimate (0.194) was lower than the sum *φ*_*z*→*w*_ +*φ*_*v*→*u*_ = 0.3, if one expects gene flow occurring in the same direction to be accumulative (see, e.g., Malinsky *et al*., 2021, Fig. 1c). The inferred *D* → *E* gene flow did not exist in the true model and reflected the fact that the triplet methods assume an inflow setup (e.g., Ji *et al*., 2023) and are unable to infer the direction of gene flow. Finally the inferred *F* → *D* gene-flow event reflected the true *v* → *u* gene flow, and to a lesser extent the *z* → *w* event as well since species *E* is an unsampled ghost species in the analysis of the *CDF* triplet. It is interesting that 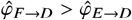.

The *x* → *y* introgression in the true model (Fig. 1**a**) was not inferred in the *f* -branch analysis of the dataset. Note that given the true model, there exists no triplet in which the *x* → *y* gene flow is ‘inflow’, as assumed by the triplet methods. While the *x* → *y* event does not fit the inflow setup, gene-flow signal in the opposite direction (*y* → *x*) was detected in the following four triplets: *EC A, FC A, E D A*, and *FD A*, with the *f*_4_ estimates to be 0.0015, 0.0017, 0.0032, and 0.0034, respectively. However, those estimates were not significant (with the *p*-value *>* 0.01) and were set to zero. Thus no gene flow, either from *x* → *y* or from *y* → *x*, was inferred in the *f* -branch analysis of the dataset (Fig. 1**c**).

#### Performance of the f -branch test

The performance of a statistical test is assessed by its false-positive rate (type-I error) and power (or 1 minus the false-negative rate). A useful test is required to have the type-I error under control (with the false positive rate to be ≤ 5% if the test is conducted at the 5% level), and then one evaluates the power of the test. If we insist on the inference of the full model of Figure 1**a**, including the correct number of gene-flow events, correct identification of populations involved in gene flow, and correct directions of gene flow, the approach achieved 0% accuracy: in none of the replicate datasets simulated in this study was the correct model ever recovered by the method.

However, there is ambiguity in deciding whether a gene-flow event inferred by the *f* -branch approach is correct or not. For example, as the triplet methods used are agnostic of the direction of gene flow, an inferred gene flow event may be considered correct if it involves the correct lineages but wrong direction. Indeed Malinsky *et al*. (2021, p.592) point out that “an *f* -branch result in itself does not indicate directionality of gene flow”, although estimates of introgression probabilities for unidirectional gene flow are reported (e.g., Fig. 3 of the same paper). The probability of introgression is meaningful only if the direction of gene flow is known. Furthermore all triplet methods used in *f* -branch to estimate the introgression probability assume inflow. Here we consider the *f* -branch estimates as meaningful, noting that the estimates may be biased if gene flow is in the opposite direction.

If the inferred *A* → *C* and *B* → *C* gene flow is considered a false positive error, the error rate was no larger than 5% at the 0.03 cut-off, and even lower at the 0.05 cut-off (table 2). If we consider those gene-flow events as correct results (reflecting the true *x* → *y* introgression in the opposite direction, Fig. 1**a**), those proportions will be the power of the test, suggesting that power was very low.

**Table 2.** False positive rates and power of *f* -branch test of gene flow in analysis of data simulated under the model of Figure 1.

| Cutoff <sup>†</sup> | Same $\theta$ | | Different $\theta$ | |
| --- | --- | --- | --- | --- |
|  | 0.03 | 0.05 | 0.03 | 0.05 |
| **False positive rates** |  |  |  |  |
| $f_C(A)$ | 0% | 0% | 0% | 0% |
| $f_C(B)$ | 0% | 0% | 1% | 0% |
| Other branches | $\leq 1\%$ | 0% | $\leq 1\%$ | 0% |
| **Power** |  |  |  |  |
| $f_U(A) (x \rightarrow y)$ | 45% | 0% | 0% | 0% |
| $f_U(B) (x \rightarrow y)$ | 44% | 0% | 0% | 0% |
| $\text{med}(f_U(A), f_U(B)) (x \rightarrow y)$ | 42% | 0% | 0% | 0% |
| $f_E(D) (z \rightarrow w)$ | 6% | 0% | 0% | 0% |
| $f_D(E) (v \rightarrow u, z \rightarrow w)$ | 100% | 100% | 100% | 100% |
| $f_D(F) (v \rightarrow u)$ | 100% | 100% | 100% | 100% |
| $\text{med}(f_D(E), f_D(F)) (u \rightarrow v)$ | 100% | 100% | 100% | 100% |

<sup>†</sup>Evidence for introgression is considered significant if the estimated introgression probability exceeds an empirical cut-off, i.e., if $f_b > 0.03$ (Malinsky *et al.*, 2018) or $f_b > 0.05$ (Malinsky *et al.*, 2021).

The method detected the *A* → *U* and *B* → *U* gene-flow events at the 0.03 cut-off in ∼ 45% of datasets simulated with the same *θ* for all populations, and in 0% of datasets simulated with different *θ*s (table 2). These gene-flow events reflected the true gene flow from *x* → *y* in the opposite direction involving the common ancestor of *A* and *B* (Fig. 1**a**). Note that the *f* -branch approach detects gene flow from donor populations that are tips but not internal nodes on the species tree. If we consider those events as correct results, both reflecting the *x* → *y* gene flow, the power for detecting the event is then 45% and 0%. If we consider the inferred *A* → *U* and *B* → *U* gene-flow events to be incorrect, those proportions will be false positive rates (table 2).

The assumption that the donor population for gene flow must be a tip branch may be problematic. One idea may be to move the gene-flow events supported by all descendant tips onto the ancestral branch, similar to the *f* -branch treatment of recipient populations. Suppose one chooses species *A*_*i*_, *B* _*j*_, *C*_*k*_ from clades *A, B, C* to form the triplet *A*_*i*_ *B* _*j*_*C*_*k*_, with estimated introgression probability *f* (*A*_*i*_, *B* _*j*_; *C*_*k*_, *O*) (Fig. 10). One may generate an estimate for the *c* → *b* gene flow involving the ancestral branches *c* and *b*,

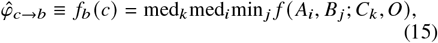

where the med_*k*_ operation is taken only if all rates involving descendent tips of branch *c* are positive. Note that in the dataset of Figure 1**c**, this approach will infer the *v* →*u* gene flow but miss the *z* → *w* gene flow.

If we apply this idea and replace the *A* → *U* and *B* → *U* gene-flow events by *y* → *x* gene flow, we will infer the correct lineages (the true event is from *x* → *y* in the opposite direction). Then power at the 0.03 cut-off will be 42% in datasets assuming the same *θ* and 0% in datasets with different *θ*s (table 2).

The *E* → *D* and *F* → *D* gene-flow events were detected in all datasets (table 2). As noted in Figure 1**c**, the *F* → *D* event reflected the true *v* → *u* event, and to a lesser extent the *z* → *w* event as well since species *E* is a ghost species in the analysis of the *CDF* triplet. The power for detecting both *E* → *D* and *F* → *D* gene-flow events was 100%.

The *D* → *E* event was detected at the 0.03 cut-off in 6% of datasets simulated with the same *θ* and in 0% of datasets with different *θ*s. This reflected the true *E → D* gene flow (Fig. 1**a**) and may be considered a correct result.

The *v* → *u* introgression in the true model involves the common ancestor of *E* and *F* as the donor population. This was not detected, because as discussed above the *f* -branch approach assumes that the donor population must be a tip branch on the species tree.

#### Performance of f -branch estimation of introgression probabilities

Estimates of introgression probabilities obtained from

*f* -branch analysis of replicate datasets simulated under the model of Figure 1**a** are summarized in Figure 11. Five gene-flow events were ever detected by the *f* -branch method (Fig. 11): *f*_*U*_ (*A*) (or *φ*_*A*→*U*_), *f*_*U*_ (*B*) (or *φ*_*B*→*U*_), *f*_*D*_ *E* (for *φ*_*E*→*D*_), *f*_*D*_ *F* (for *φ*_*F*→*D*_), and *f*_*E*_ *D* (for *φ*_*D* →*E*_). The first two events, *A* → *U* and *B* → *U*, reflect the *x* → *y* gene flow in the true model although the true direction is the opposite and the true lineage is a parental lineage (Fig. 1**a**). In general the estimated rate for those two events were low, except for datasets simulated under the same *θ* (Fig. 11). The assumed wrong direction of gene flow may partly account for the low estimates, and the truncation used by Dsuite, which sets the *f*_4_ estimate of *φ* for any triplet to zero if the associated *D*-statistic is not significant (Malinsky *et al*., 2021), may be another factor.

**Fig. 11.**
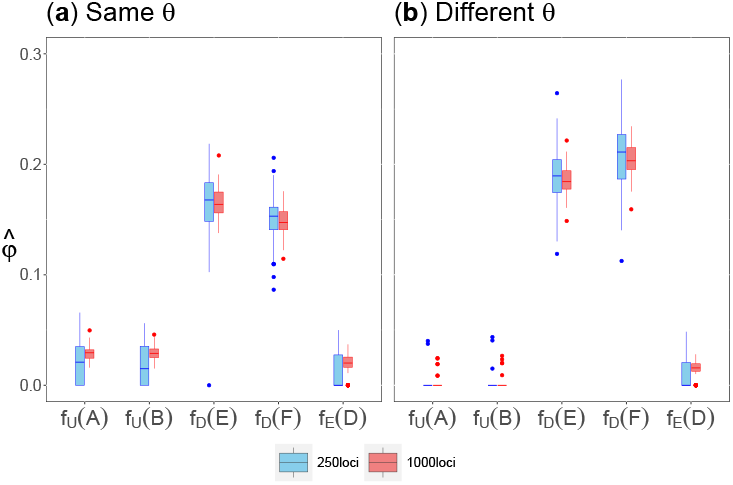
Boxplots of estimates of introgression probability (*φ*) from the *f* -branch analyses of data simulated under the six-species model of Figure 1**a**, with either *L* = 250 or 1000 loci in each dataset. Only five introgression events were ever found to be significant (with FWER *<* 0.001) in the data simulated in this study.

The other three events, *E* → *D, F* → *D*, and *D* → *E*, all reflect the true gene flow from *E* → *D* (that is, *z* → *w* and also *v* → *u*; Fig. 1**a**).

We consider the *F* → *D* introgression to be a correct result as well; the detected signal may reflect both the *V* → *D* and *E* → *D* gene flow involving the sister species *E* of *F*. If the introgression probabilities are additive and independent of the direction, one would expect *φ*_*E* →*D*_ to be close to *φ*_*v*_ + *φ*_*w*_ = 0.2 + 0.1 = 0.3. Estimates from *f* -branch were around 0.1, much lower. The *f*_*D*_ *(F*) estimates were smaller than and close to 0.1, also much lower than *φ*_*v*_ = 0.2.

#### Analysis of the six-species data using BPP

We analyzed the data simulated under the six-species model with three introgression events (Fig. 1**a**) using BPP, specifying the correct MSC-I model. The posterior means and the 95% HPD CIs for parameters in the true model were well behaved (Fig. S4). The CIs were narrower in large datasets of *L* = 1000 loci than in small datasets (*L* = 250). The estimates were far more precise with much narrow CIs in datasets simulated under the same *θ* for all populations than for datasets with different *θ*s: if all species on the phylogeny have the same population size (*θ*) there is a large benefit in parameter estimation in enforcing the assumption.

We calculated the Bayes factor in support of gene flow (*B*_10_) using the S-D density ratio (Ji *et al*., 2023). In every dataset, the Bayes factor for testing each of the three introgression events was ∞ (except for one dataset of 250 loci), strongly supporting gene flow.

We also analyzed the *CDE* triplet data (including the outgroup *O*). We used the data simulated under the six-species model with *L* = 1000 loci, to fit four variants of the MSC-I model: (i) the true model with two introgression events (*v* → *u, z* → *w*; Fig. 1**b**), (ii) bidirectional introgression (BDI, *z* ↔ *w*), (iii) unidirectional introgression (UDI) from *E* → *D* (or *z* → *w*), and (iv) UDI in the wrong direction from *D* → *E* (or *w* → *z*). Estimates of the introgression probabilities are summarized in table 3.

**Table 3.**
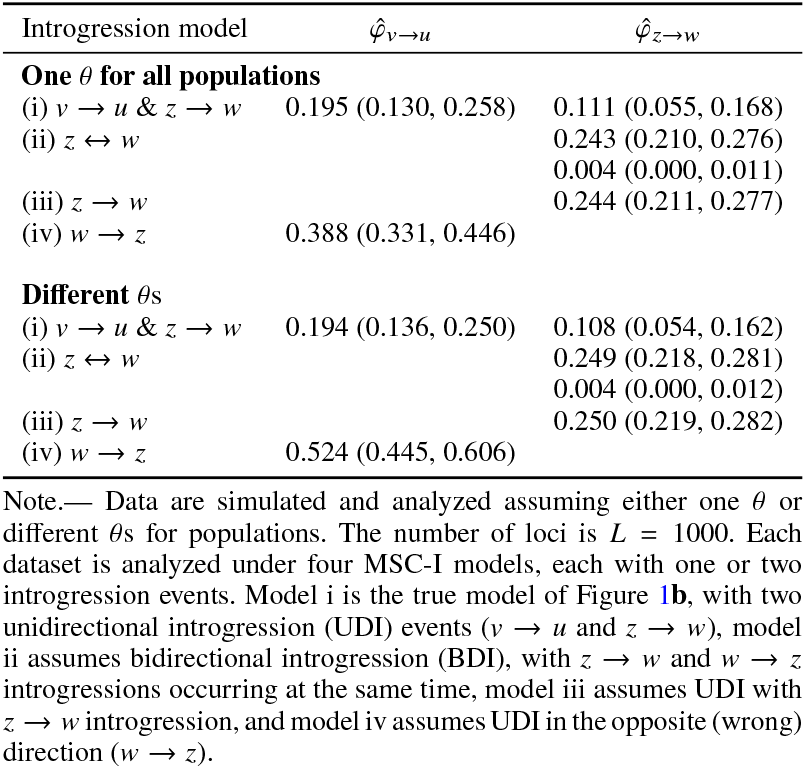
Average posterior means and 95% HPD CIs for introgression probabilities (*φ*) in BPP analyses of the *CDE* triplet data (including outgroup *O*) simulated under the six-species model of Figure 1**a**.

Model i, with two UDI gene-flow events (*u* →*v* and *z* → *w*), constitutes the true model for those data. Under this model the estimates were highly accurate. For example the average posterior means for *φ*_*v* → *u*_ (with true value 0.2) were 0.195 for datasets with the same *θ* and 0.194 for datasets with different *θ*s, while the corresponding estimates for *φ*_*z* → *w*_ (with true value 0.1) were 0.111 and 0.108, respectively. The estimates were slightly less precise than those from the full data for all six species (Fig. S4) due to reduced information content. Model ii (BDI) assumes introgression in both directions (*z* ↔ *w*): the estimates were 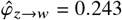 or 0.249 in the correct direction and 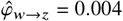 in the wrong direction. The estimates were largely the same as those from model iii assuming the *z* → *w* introgression only. This estimated rate was somewhat accumulative (compare 0.243 with the true rate *φ*_*v*→*u*_ + *φ*_*w*→*z*_ = 0.2 0.1 = 0.3. Model iv assumes one introgression event in the wrong direction, and the estimated rate was even higher than the true rate in the correct direction (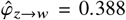 for data of the same *θ* and 0.524 for data of different *θ*s, compared with *φ*_*v*→*u*_ +*φ*_*w*→*z*_ = 0.3).

Bayes factors strongly supported gene flow in each of the four models of table 3 except the *w* → *z* gene flow in model ii (BDI), in which the gene-flow event was strongly rejected (with *B*_10_ *<* 0.01). Note that in model iv with the only assumed gene flow in the wrong direction, gene flow was also strongly supported, consistent with the large estimate of the rate.

Those results are highly consistent with Thawornwattana *et al*. (2023). In particular, if introgression is assumed to occur in the wrong direction, the test of gene flow will often be significant and the estimated rate may even be greater than the true rate in the opposite direction. However, if bidirectional introgression is assumed, gene flow in the correct direction will be detected and the nonexistent gene flow in the wrong direction will be rejected (Thawornwattana *et al*., 2023).

## Discussion

### Impacts of incomplete taxon sampling on inference of gene flow

In this study, we have examined mainly two effects of incomplete taxon sampling on inference of gene flow using genomic data: (i) the impact on ghost lineages that are donors of gene flow into modern species (i.e., ghost introgression) and (ii) the impact of changing population sizes when some species are unsampled or excluded in triplet analyses.

As demonstrated by Tricou *et al*. (2022) and Pang and Zhang (2024), existence of ghost donor lineages can mislead methods such as *ABBA*-*BABA* and HyDe to infer gene flow involving incorrect lineages. Our simulation suggests that a Bayesian testing approach (Fig. 2) can distinguish different models of gene flow such as ghost introgression, inflow, and outflow, with excellent sensitivity (high power) and specificity (low false positives) (table 1). It may be worth investigating whether the BPP analysis of triplets, as examined in table 1, can be used to replace triplet methods used in the *f* -branch approach.

We note that our study of ghost introgression has very limited scope. There exist many scenarios in terms of the size and shape of the species phylogeny, the number of gene-flow events, the lineages involved (ancestral versus modern), the mode of gene flow (discrete versus continuous), and so on. We leave it to the future to investigate the problem more systematically.

Incomplete taxon sampling may also cause multiple branches on the phylogeny corresponding to species with different population sizes to be merged into one species, with one population size assumed in the model. A similar situation arises when the population size of the same species changes over time (as in Fig. 3**a**). We find that the introgression probability (*φ*) is well-estimated if incomplete taxon sampling causes multiple branches on the species tree to be merged into one branch representing the recipient population of gene flow, even though the population size for the recipient population may involve large errors or biases (Figs. 6&8). This appears to be due to the fact that the parameter that influences gene genealogies and that can be reliably estimated from genomic data is the expected proportion of immigrants in the recipient population at the time of hybridization/introgression (*φ*), rather than the expected number of immigrants. We also find that estimates of the rate of gene flow by both Bayesian and summary methods may be seriously biased if different species have different population sizes but genomic data are analyzed assuming one population size for all species (Figs. S3&9).

Incomplete taxon sampling may also cause multiple gene-flow events, which occur at different time points in different directions, to be merged onto the same branches in the tree for the species subset (compare the *v* → *u* and *z* → *w* gene-flow events in the full phylogeny of Fig. 1**a** with the events in the triplet tree of Fig. 1**b**), or may cause gene flow between non-sister species to become gene flow between sister species. Because commonly used summary triplet methods cannot identify gene flow between sister lineages and assume only one gene-flow event between nonsister lineages, it may be impossible for those methods to recover the correct gene-flow events in such scenarios. There is a need to improve triplet methods, for example, to have the power to infer gene flow between sister lineages.

### The f -branch approach to inferring gene flow

The *f* -branch approach is an exploratory method for integrating triplet estimates of introgression probabilities when data from many species are analyzed using summary triplet methods (Malinsky *et al*., 2018, 2021). Our evaluation here suggests that if there likely exist multiple introgression events involving ancestral branches, interpretation of the *f* -branch results may not be straightforward. Here we provide a summary of our findings as well as the simulation results of Malinsky *et al*. (2018, 2021), which may aid the interpretation of results obtained from an *f* -branch analysis.

First, note that triplet methods used in the *f* -branch approach, such as the *D*-statistic, *f*_4_ (Green *et al*., 2010, SOM18), *f*_*d*_ (Martin *et al*., 2015, Fig. 1) and HyDe(Blischak *et al*., 2018, Fig. 1), are unable to identify gene flow between sister lineages. It appears that the *f* -branch approach should be able to infer gene flow between sister lineages as well if a triplet method that can identify gene flow between sister lineages is used instead.

Second, the *f* -branch approach provides estimates of the introgression probability (*φ*) by summarizing estimates generated by triplet methods such as *f*_4_ (Green *et al*., 2010, SOM18), *f*_*d*_ (Martin *et al*., 2015, Fig. 1) and HyDe(Blischak *et al*., 2018, Fig. 1), all of which assume inflow (i.e., from branch *C* to *B* rather than from *B* to *C* in Fig. 10). However, outflow may also show up as signals of gene flow in such analyses, as gene flow in opposite directions generate similar signals in sequence data, such as a reduction in genetic distance between the lineages involved. Estimates of introgression probability by summary methods (and thus by *f* -branch) may be biased if gene flow is in the outflow direction or if gene flow occurs in both directions (see, e.g., Ji *et al*., 2023; Thawornwattana *et al*., 2023). Note that introgressions in opposite directions between the same two lineages may neither cancel out nor be additive (Thawornwattana *et al*., 2023). The pattern may be complex, depending on the population sizes of the donor and recipient lineages and the rates in the two directions (Thawornwattana *et al*., 2023).

Third, commonly used triplet methods for estimating the introgression probability assume one population size (*θ*) for all species on the phylogeny, and estimates may involve large biases if population size differs among species on the phylogeny or change over time (Fig. 9).

Fourth, the *f* -branch approach identifies only gene flow from a tip branch on the species tree; in other words, the donor population must be an extant lineage. If both the donor and the recipient populations correspond to ancestral branches on the species tree (e.g., the *x*→ *y* and *v* → *u* gene-flow events in the model tree of Fig. 1**a**), the method will have no chance of inferring the correct gene flow. Furthermore, if the donor population is ancestral to the *C* clade, the single gene-flow event may show up in many *f* -branch tests (Fig. 10).

In summary, most limitations of the *f* -branch approach discussed here concern the triplet methods used. Improvements in triplet methods may be expected to lead to improvements in the *f* -branch analysis as well. Overall the *f* -branch approach is effective in generating plausible gene-flow scenarios when there are overwhelmingly many triplets with significant gene flow. We suggest that *f* -branch is a useful tool in analysis of genomic datasets to generate hypotheses of gene flow that may be tested using more rigorous methods.

## Supplementary Material and Data Availability

Supplemental information is available at https://doi.org/10.5061/dryad.cvdncjtgt.

