## Supplemental Figures for "The impact of incomplete taxon sampling on inference of gene flow by Bayesian and summary methods using genomic sequence data"

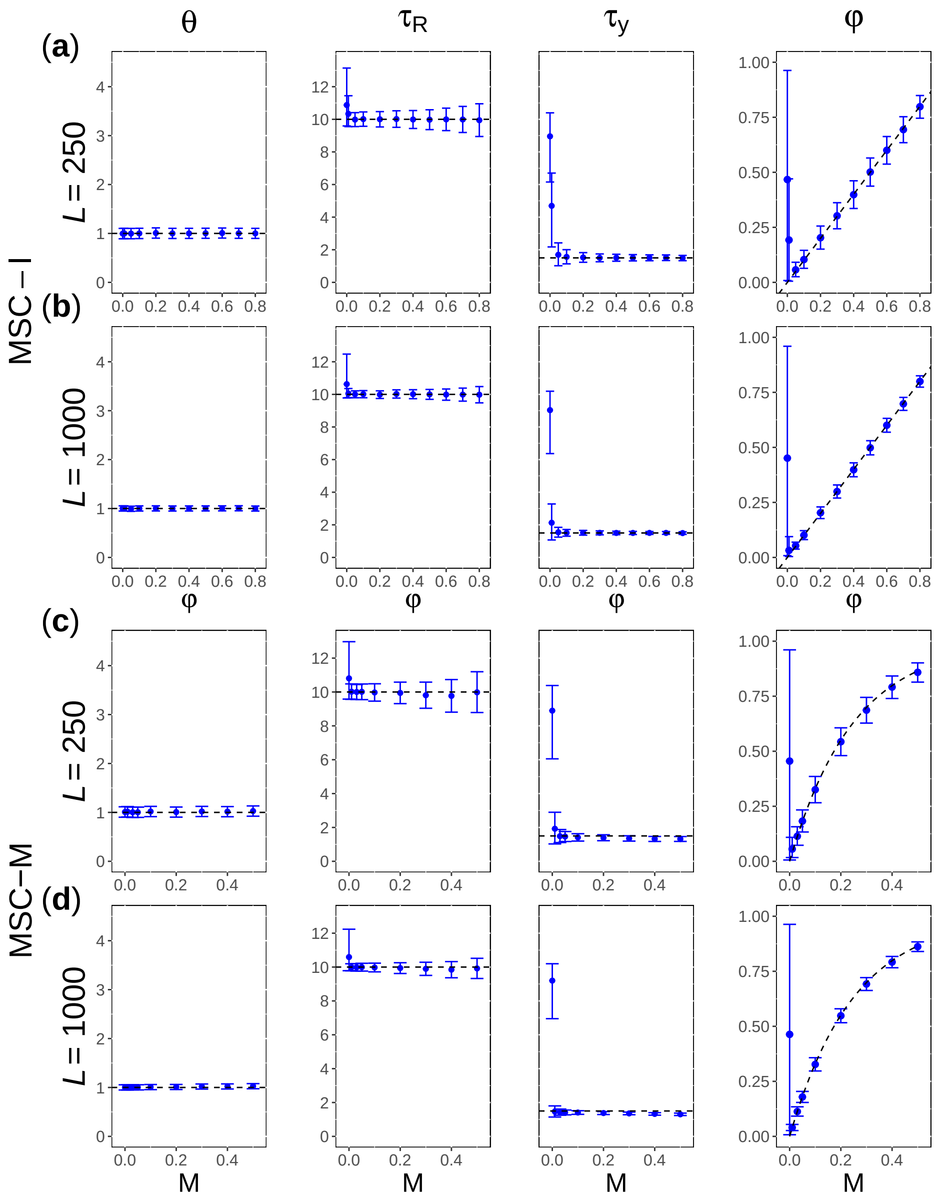


**Fig. S1**: [2s-same-same] (**a**, **b**) Average posterior means and the 95% HPD CIs of parameters when the data are simulated and analyzed under the MSC-I model (Fig. 4**a**) assuming the same $\theta$ for all species, plotted against the introgression probability ($\varphi$). (**c**, **d**) Parameter estimates under MSC-I when data are simulated under MSC-M (Fig. 4**b**), plotted against the migration rate ($M$). Estimates of $\theta$ and $\tau$ are mutipled by 1000. Black dashed lines represent the true value in **a&b** or expected value ($\varphi_0$) in **c&d**.

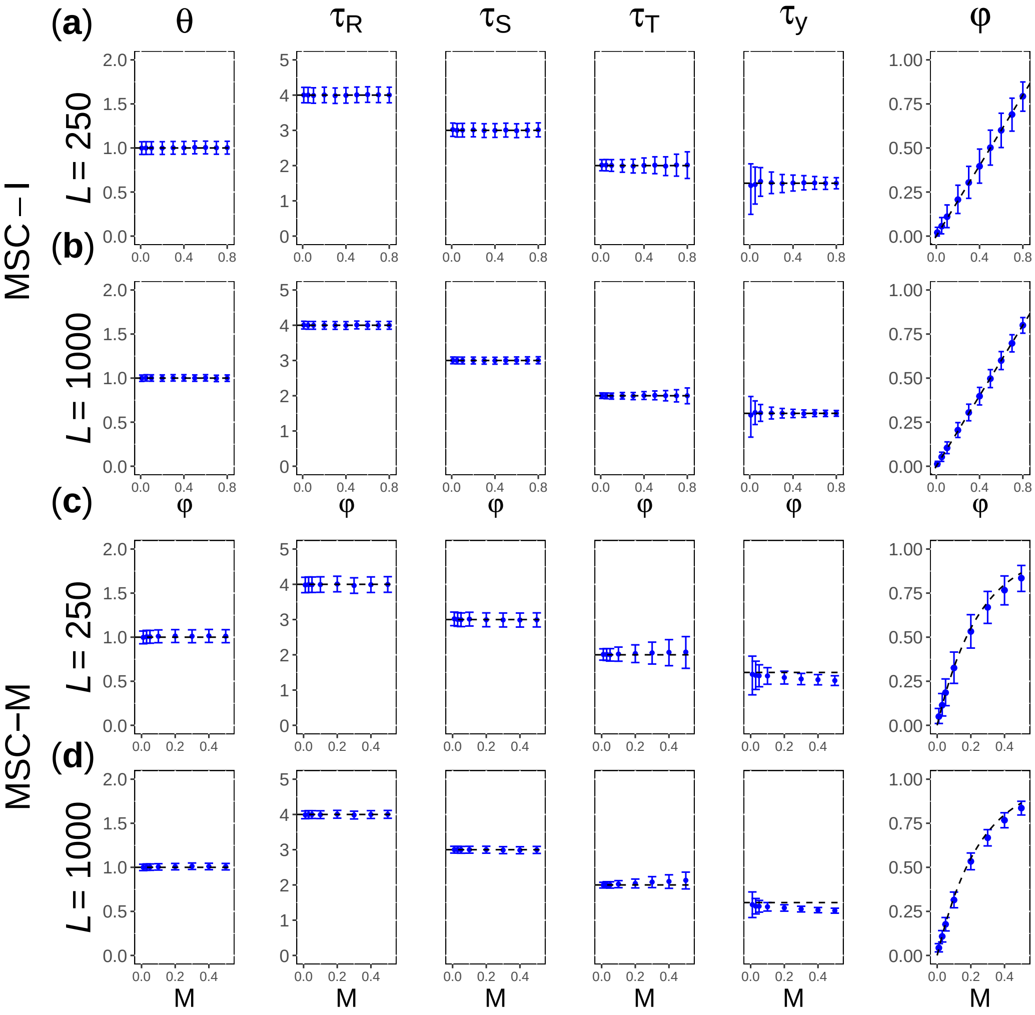


**Fig. S2**: [4s-same-same] (**a**, **b**) Average posterior means and the 95% HPD CIs of parameters from data simulated and analyzed under the MSC-I model (Fig. 5**a**) assuming the same $\theta$ for all populations, plotted against the introgression probability ($\varphi$). (**c**, **d**) Parameter estimates under MSC-I from data simulated under MSC-M (Fig. 5**b**), plotted against the migration rate ($M$). Dashed lines represent true parameter values except that in **c&d** for $\tau_y$ and $\varphi$, they represent $(\tau_T + \tau_U)/2$ and $\varphi_0$ (eq. 13), respectively.

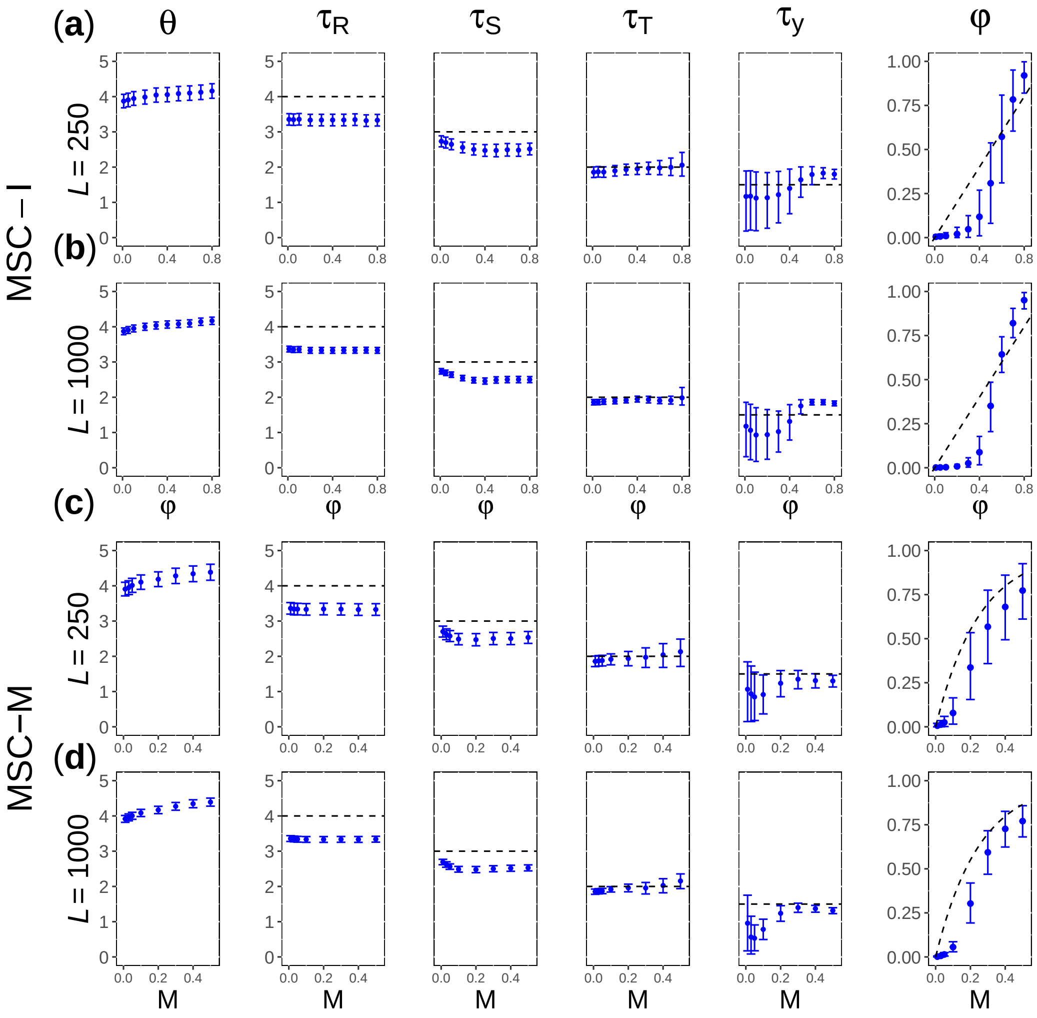


**Fig. S3**: [4s-diff-same] (**a**, **b**) Average posterior means and the 95% HPD CIs of parameters under MSC-I (Fig. 5**a**) from data simulated under the MSC-I model (Fig. 5**a**) with different $\theta$s for species, but assuming the same $\theta$ for all populations in data analysis. (**c**, **d**) Estimates under MSC-I when the data are simulated under the MSC-M model (Fig. 5**b**)

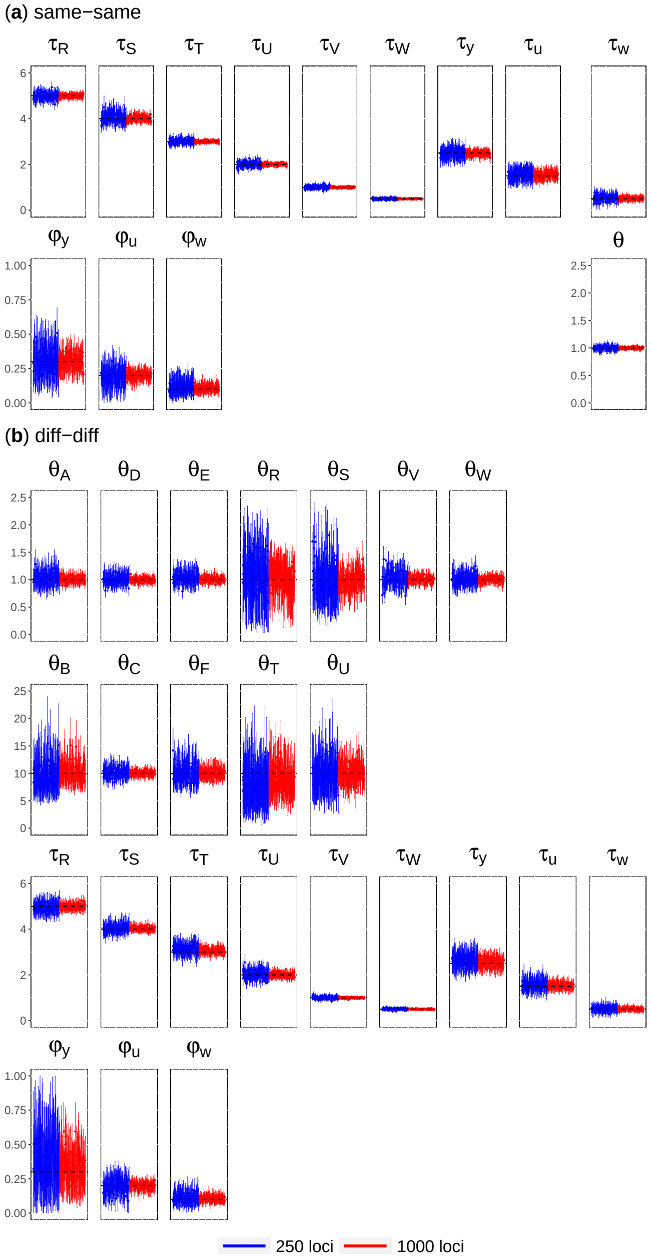


**Fig. S4**: Posterior means and the 95% HPD CIs for parameters when replicate datasets are simulated and analyzed under the MSC-I model of Figure 1**a** assuming either (**a**) the same $\theta$ for all populations or (**b**) different $\theta$ for populations. Black dashed lines represent the true values.
